# An RNA seq-based reference landscape of human normal and neoplastic brain

**DOI:** 10.1101/2023.01.03.522658

**Authors:** Sonali Arora, Frank Szulzewsky, Matt Jensen, Nicholas Nuechterlein, Siobhan S Pattwell, Eric C Holland

## Abstract

In order to better understand the relationship between normal and neoplastic brain, we combined five publicly available large-scale datasets, correcting for batch effects and applying Uniform Manifold Approximation and Projection (UMAP) to RNA-seq data. We assembled a reference Brain-UMAP including 702 adult gliomas, 802 pediatric tumors and 1409 healthy normal brain samples, which can be utilized to investigate the wealth of information obtained from combining several publicly available datasets to study a single organ site. Normal brain regions and tumor types create distinct clusters and because the landscape is generated by RNA seq, comparative gene expression profiles and gene ontology patterns are readily evident. To our knowledge, this is the first meta-analysis that allows for comparison of gene expression and pathways of interest across adult gliomas, pediatric brain tumors, and normal brain regions. We provide access to this resource via the open source, interactive online tool Oncoscape, where the scientific community can readily visualize clinical metadata, gene expression patterns, gene fusions, mutations, and copy number patterns for individual genes and pathway over this reference landscape.

## Introduction

Over the past several decades the scientific community has characterized and cataloged individual genes for their function and involvement in development and disease. More recently, several single-cell atlases from different tissues and organ sites for various species (for example human and mouse) have been created that provide deep insights into the relationships between different cell types in development and in adult tissues. In this study, we have blended publicly available RNA seq datasets of normal and neoplastic brain to derive similar insights into the relationship between various central nervous system (CNS) tumors and between neoplastic vs. normal brain. Using this reference landscape, expression of specific genes and gene ontology groups can be compared across all tumor types and normal brain regions.

A variety of omics approaches have been employed to characterize both tumor and healthy tissue at the molecular level by various large-scale international initiatives. The Cancer Genome Atlas (TCGA)^1^ contains data from across 33 cancer types, including uniformly processed cancer genomic data (whole transcriptome RNA-Seq), microarray, gene fusions, gene mutations and copy number calls) from 702 glioma patients and 5 matched normal patients. The Chinese Glioma Genome Atlas (CGGA)^2^ contains RNA-seq, whole genome sequencing, DNA methylation, microarray data from over 2000 brain tumor samples. The Children’s Brain Tumor Tissue Consortium (CBTTC)^3^ contains whole genome sequencing and RNA-seq data across 23 different pediatric tumors. The Genotype Tissue Expression Project (GTEx)^4^, contains genomic data from 54 non-diseased tissue sites across nearly 1000 individuals, including 1409 brain tissue samples from 13 GTEx defined brain regions.

Here, we present a visual integration approach for analyzing multiple diverse molecular datasets combining large numbers of samples across different brain regions and brain tumor subtypes. We combined transcriptional data from three different datasets for adult glioma, pediatric tumors and healthy normal brain regions, corrected for batch effects, and used a dimension reduction technique to construct a UMAP to find meaningful patterns in this pooled multi-disease dataset.

This comparative study can help researchers and clinicians visualize similarities and differences in patient cohorts, study and compare alterations in gene expression, signaling pathways, gene fusions, copy number profiles and mutation calls across multiple tumor types. By adding normal healthy brain tissues from GTEx to our reference landscape, we also allow for comparisons between healthy and neoplastic states. Visualizing these similarities and differences in an opensource, interactive website, Oncoscape ^5^(https://oncoscape.sttrcancer.org/#project_bulkrnaseqbrainumap), can aid in analysis for translational research.

## Results

### Constructing the Brain-UMAP / Clustering of gene expression data identifies diverse disease types

To characterize and better understand the molecular intricacies of brain tumors, we downloaded uniformly processed RNA-seq abundances values from recount-brain, a curated repository for human brain RNA-seq datasets, for three different uniformly processed datasets - 702 adult glioma samples from TCGA^1^, 270 adult glioma samples from CGGA^5,6^, 1409 healthy normal brain samples from GTEx^4^ across 12 GTEx-defined brain regions (**Supplementary Table 1a**). Retrieving data from recount^7^ ensured that consistent bioinformatic pipelines were used for these three datasets thus resulting in no batch effects between the three datasets.

The most common solid tumors in children are brain tumors with approximately 1.15 to 5.14 cases per 100,000 children in the United States alone^8^. To adequately represent a wide range of CNS tumors in our reference landscape, we additionally included 802 pediatric tumor samples (**Supplementary Table 1b)** from the Children Brain Tumor Network (CBTN)^3^. **Fig. 1a** represents an overview of data sources (details in **Supplementary Table 1c**)

**Fig1.**
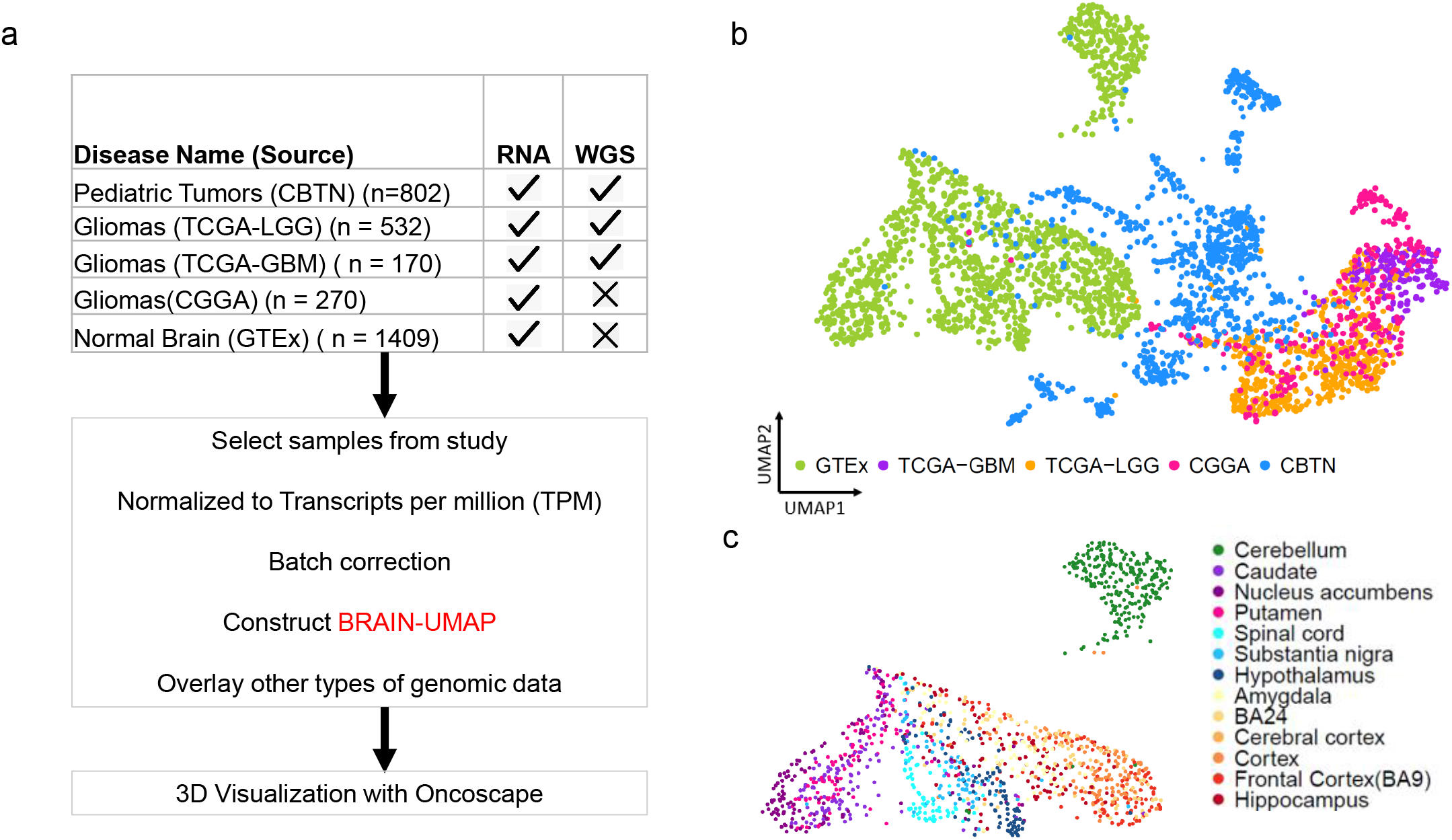
Overview of data analyzed (a) showing datasets used, batch correction and construction of Brain-UMAP. (b) UMAP of complete dataset including adult gliomas, pediatric tumors and GTEx-defined normal brain. (c) UMAP showing unique clustering of GTEx-defined brain-regions

Gene expression data from each of the datasets was converted to units of Transcripts per million (TPM) to avoid inter-pipeline difference and were limited to a common set of 19142 protein-coding genes^9^. While both CBTN and recount used different bioinformatic pipelines (**Supplementary Table 1c**), in order to ensure that there were no batch effects we used combat^10^ method from the R package “sva” to remove unwanted variation in our combined dataset.

We explored three different dimension reduction techniques (Principal Component Analysis (PCA), t-distributed Stochastic Neighbor Embedding (tSNE) and uniform manifold approximation and projection (UMAP) for data visualization. We chose UMAP to build a Brain-UMAP (**Fig. 1b**) on batch corrected TPM integrated counts, as UMAP segregated the mini clusters well and was very effective in visualizing clusters and their relative proximities (**Supplementary Fig 1a**).

While 2-dimenisonal representation of the data is helpful, we also provide a 3-dimensional representation of the data on the interactive web-based platform Oncoscape^5^) where users can easily toggle among and compare different patient groups, while using a suite of interoperable tools.

At a first glance, distinct clustering of samples is observed. The adult glioma samples from the different datasets, TCGA-LGG, TCGA-GBM and CGGA, co-localized closely within the same region of the UMAP, whereas the pediatric tumor samples clustered between the GTEx healthy normal brain and the adult glioma samples. (**Fig. 1b**).

### Distinct gene expression profiles in normal human brain

The 1409 normal brain samples segregated into two distinct clusters of multiple supratentorial regions and cerebellum, as we have previously shown^11^ (**Fig. 1c**). The supratentorial regions further revealed three anatomically distinct regions for basal ganglia (caudate, nucleus accumbens, putamen), cortex (amygdala, Brodmann Area 24, cerebral cortex), and the midline structures (spinal cord, substantia nigra and hypothalamus). We confirmed that different classifiers such as postmortem interval (PMI), age (years), sex, Hardy score and type of sample preparation (**Supplementary Fig. 1b**) were not associated with distinct clustering patterns, suggesting that the sample clusters we observe are based on actual biological differences between these brain regions rather than sample preparation parameters. However, we observed that a few samples with much lower RNA integrity number (RIN) compared to all the other samples from different brain regions converged at a point (**Supplementary Fig. 1b**).

### Clustering of transcriptomic glioma datasets reveals distinct glioma subtype

Within the adult glioma clusters, we observe that while the samples from the two TCGA datasets cluster together, there is a more continuous pattern when looking at gene expression profiles for samples from TCGA-GBM and TCGA-LGG (**Fig. 2a**). This contrasts with the distinct clusters observed by analyzing whole exome single nucleotide alterations (SNAs) and whole genome copy number alterations (CNAs) from the same patients, as shown by Bolouri^12^ et al. In line with previous reports, we observed that the age of the patients at diagnosis for the TCGA-GBM and TCGA-LGG samples revealed a sharp gradient illustrating the known correlation between age and outcome (**Fig. 2b**). By contrast, patient gender was not associated with any specific clusters (**Supplementary Fig. 2a**). Regarding chromosomal alterations, tumors with a gain of chromosome 7 and hemizygous deletion of chromosome 10 (**Fig. 2c**) or co-gain of chromosome 19 and 20 (**Supplementary Fig 2b**) co-localized in the top area of the UMAP containing the TCGA-GBM samples (**Fig. 2c**). Tumors exhibiting a co-deletion of chromosome 1p and 19q and mutation in isocitrate dehydrogenase 1 and 2 genes (IDH-mut) (**Fig. 2d**) were concentrated in the lower half of the UMAP containing the TCGA-LGG samples. Tumors containing mutation in IDH1 (**Fig. 2e**), TP53 (**Fig. 2f**) and ATRX (**Fig. 2g**) were also found to be concentrated and clustered together in specific regions of the adult glioma UMAP. Using the supervised DNA methylation classification, transcriptional subtype, MGMT promoter status and TERT promoter status (**Supplementary Fig 2c-f**) to color the adult gliomas from TCGA-LGG and TCGA-GBM also revealed distinct patterns across the adult glioma landscape. Selecting for common glioma mutations and copy number alterations clearly shows three distinct subtypes of glioma - IDHmut-1p19q co-deleted oligodendrogliomas, the IDH mutated astrocytomas with p53 and ATRX mutations, and the wild-type IDH (IDH-wt) glioblastomas with gain of chromosome 7 and loss of chromosome 10 molecular GBM. (**Fig. 2h**).

**Fig 2.**
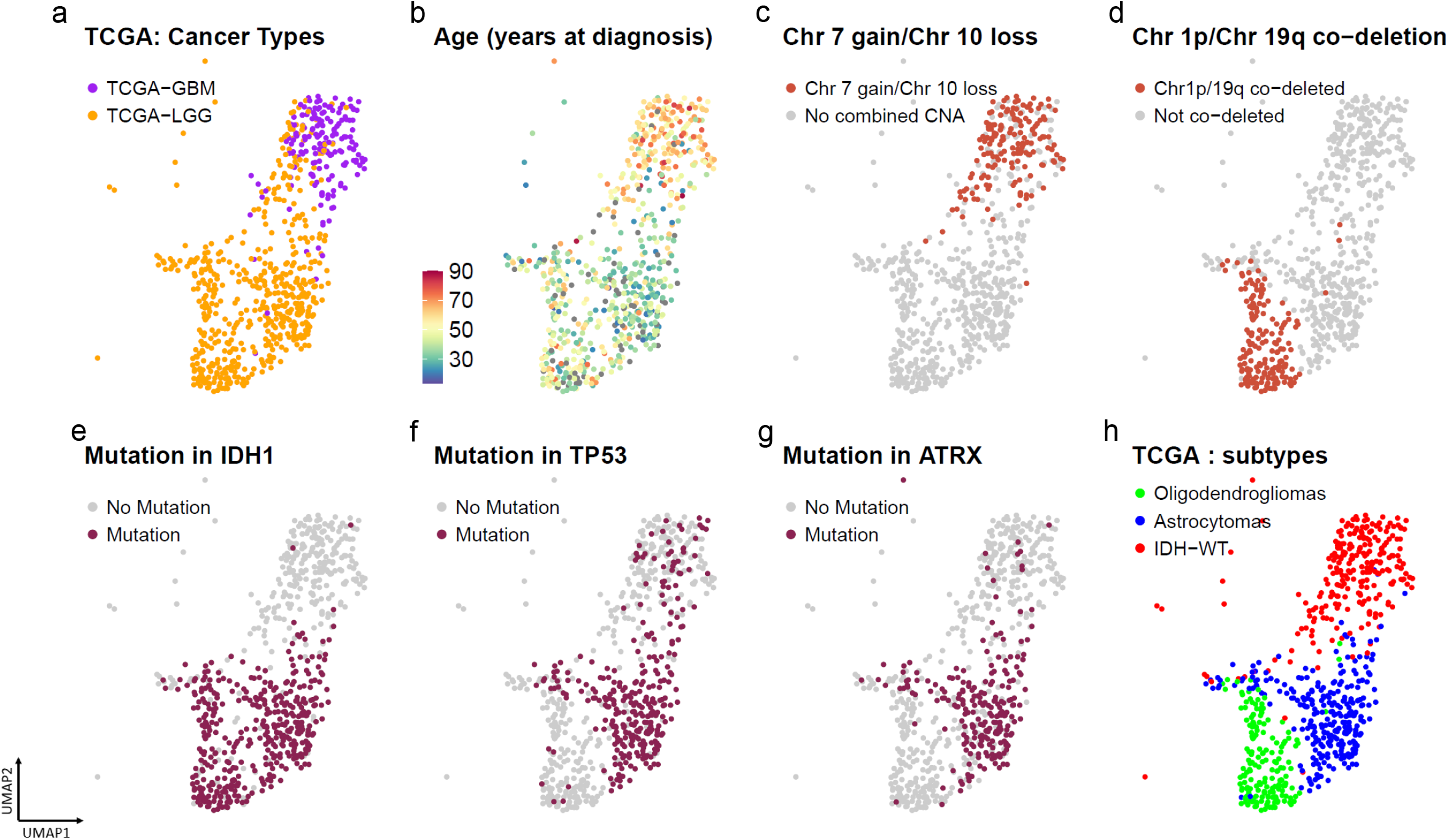
UMAP for adult glioma showing patients colored in by (a) TCGA-GBM and TCGA-LGG patients (b) age at diagnosis (c) Chr 7 gain/ Chr10 loss in patients. (d) Chr 1p/19q co-deletion status in patients (e) IDH1 mutation (f) TP53 mutation (g) ATRX mutation. (h) UMAP identifying the 3 distinct adult glioma subtypes – IDH wildtype, Astrocytoma and Oligodendrogliomas.

### Adult glioma subtypes from TCGA and CGGA show similar gene expression profiles

We next assessed if glioma samples of similar molecular subtypes from the TCGA and CGGA datasets exerted similar gene expression profiles and co-localized in their respective clusters. Similar to the TCGA samples, using grade, IDH mutation status and chromosome 1p 19q codeletion status for the CGGA samples, three distinct subtypes of adult glioma from the CGGA samples were observed. Interestingly, we observed that the oligodendrogliomas, astrocytomas and IDH-wt tumors from both CGGA and TCGA colocalized (**Fig. 3a-f**). Separated from the large cluster containing adult glioma subtypes, there were two small clusters of gliomas from the CGGA dataset. Based on their grade and IDH mutation status, one cluster consisted of a mix of IDH mutant grade 2 and grade 3 oligodendrogliomas while the second cluster consisted of grade 4 IDH-wt glioblastomas. For the remainder of the paper, we will refer to the adult glioma datasets from TCGA and CGGA by their molecular subtypes (oligodendrogliomas, astrocytomas and IDH-wt glioblastomas).

**Fig 3.**
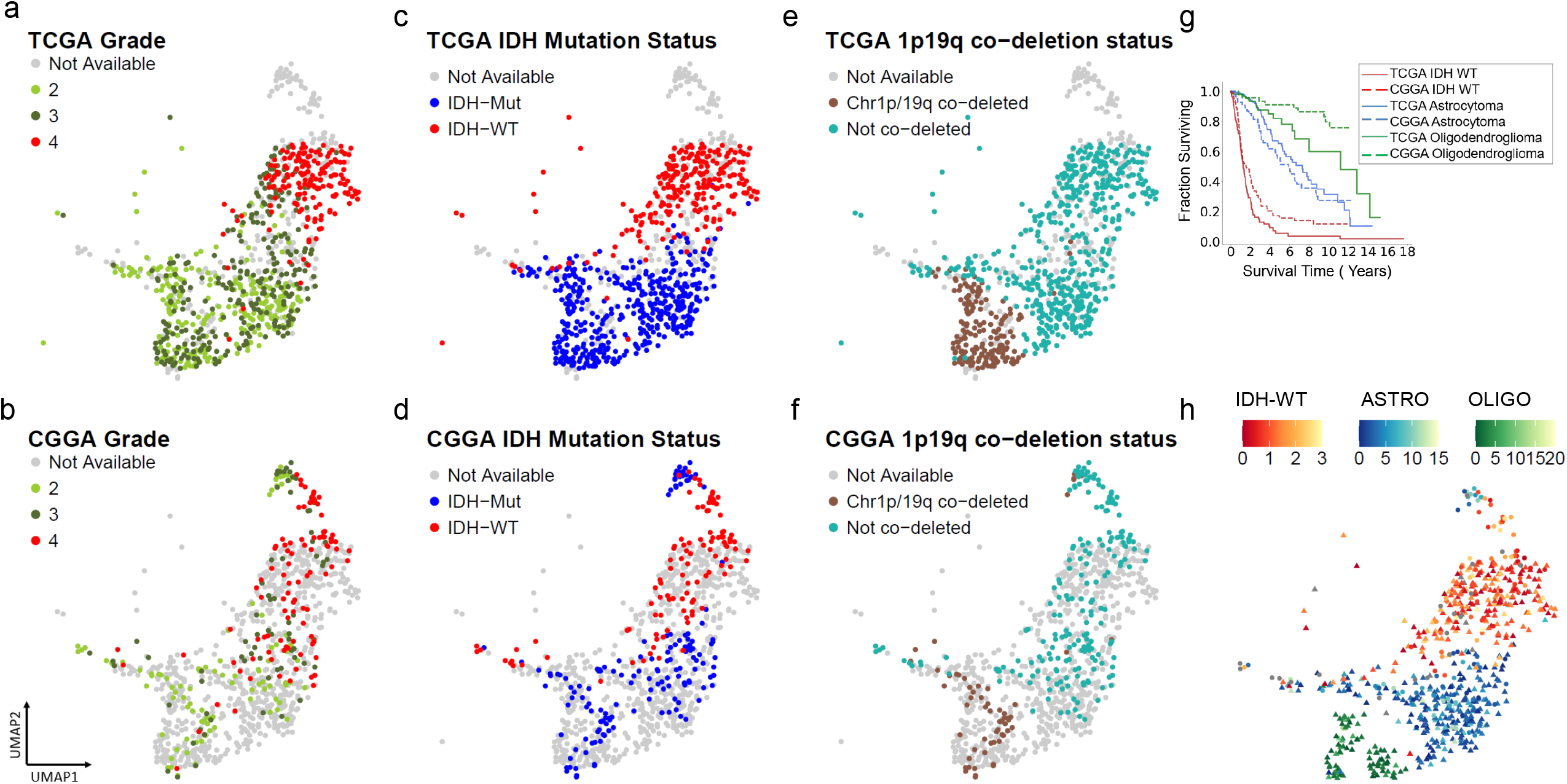
Co-localization of adult gliomas from two publicly available datasets TCGA and CGGA. Top and bottom panel shows (a) grade, (b) IDH mutation status and (c) 1p19q co-deletion status for adult gliomas from TCGA colored in, and adult gliomas from CGGA greyed out and vice versa. (d) Survival analysis for adult glioma subtypes IDH wildtype(red), Astrocytoma (blue) and Oligodendroglioma (green) from TCGA and CGGA shown in solid and dotted lines respectively. (e) Prediction of survival time (in years) using a nearest neighbor approach shows a gradient for adult gliomas.

We analyzed the survival for each subtype for all three glioma subtypes and observed similar survival rates between TCGA and CGGA datasets for the respective subtypes. (**Fig. 3g**). We then used a nearest neighbor approach to predict the survival for different UMAP subregions. Survival was predicted for samples with the median survival of its nearest neighbors, present in close proximity on the UMAP landscape (**Fig. 3h**).

We observed a small number of glioma samples formed an isthmus connecting to the normal brain samples. These samples were characterized by a low amount of copy number alterations and were a mix of IDH-wt glioblastomas, IDH-mut oligodendrogliomas or astrocytoma. These tumors were characterized by longer survival compared to other gliomas of their respective molecular subtype. It is conceivable that in some samples this region may represent either early forms of gliomas or a mix of glioma and normal brain.

Taken together, our results show that the transcriptomic data from different datasets can be combined to generate a population of adult gliomas, creating a landscape where the location of specific tumor sample can be predictive of subtype and outcome.

### Pediatric Tumors cluster by disease type

We added the 802 pediatric tumors from the CBTN dataset to our analysis to compare their gene expression patterns to both normal brain and adult glioma samples (**Fig. 4a-b**). We observed the formation of distinct subclusters for several tumor types that correlated with established molecular subgroups. As an example, we observed that medulloblastoma samples were split into three distinct clusters that correlated with known Medulloblastoma subtypes^13^ (Wnt, Sonic hedgehog (SHH) and groups 3,4, **Supplementary Fig. 4a**). Similarly, Ependymomas (EPN) samples formed several clusters that correlated with the anatomic tumor location (supratentorial (ST)-EPN, spinal-EPN, and posterior fossa (PF)-EPN) ^13^(**Fig. 4a, Supplementary Fig. 4b**). Pediatric pilocytic astrocytomas (PAs) and pediatric low-grade gliomas clustered closely together suggesting that they exert similar gene expression patterns. The schwannomas separated from the pediatric tumors and were located near the neurofibromas, which were localized adjacent to the malignant peripheral nerve sheath tumors (MPNST). The subependymal giant cell astrocytoma (SEGA) form a tight cluster as do the meningiomas. Interestingly, we observed that neurocytomas, DNET, ganglioglioma clustered near normal brain samples, specifically the hypothalamus and amygdala samples. Building a UMAP with just normal brain samples and the pediatric tumors, showed similar clustering profile (**Supplementary Fig. 4c**). Taken together, these results suggest that pediatric tumors cluster by disease type and also form subclusters made by subtypes in the case of medulloblastomas and ependymomas.

**Fig 4.**
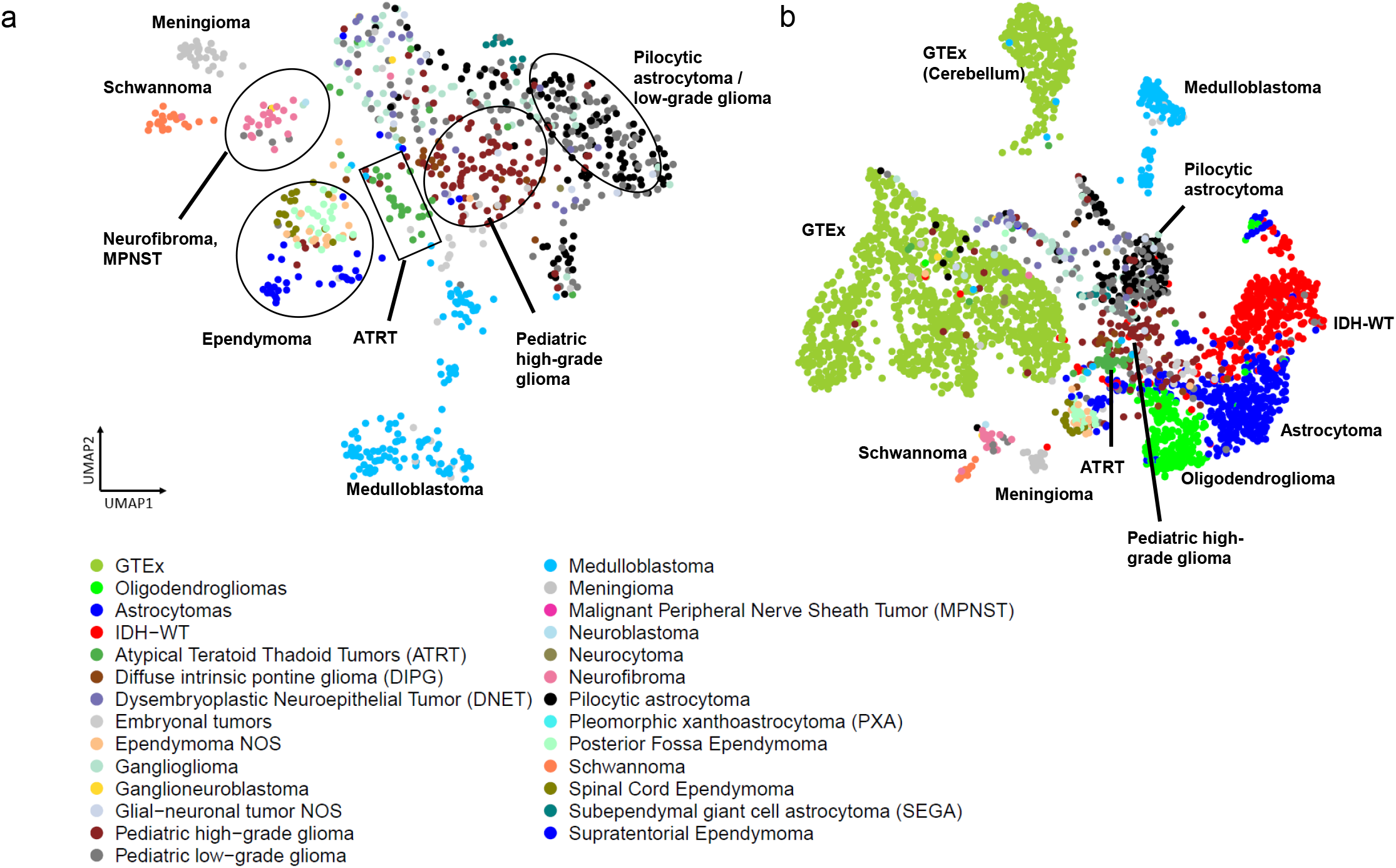
(a) UMAP of pediatric tumors (b) Updated coloring of the Brain-UMAP showing pediatric tumors and three subtypes for the adult gliomas.

### Using the reference landscape to understand pathway regulation

Alterations in signaling pathways are a hallmark of cancer and understanding the extent to which these pathways are dysregulated in tumor samples compared to healthy normal brain can help inform researchers about the underlying mechanisms of different cancer types.

Bulk gene expression from adult glioma, pediatric brain tumors and healthy brain samples was subjected to a Gene Set variation Analysis (GSVA) and the gene set variation scores for each pathway were used to color in the Brain-UMAP. A score closer to 1 represented up-regulation of pathway in the given samples, whereas a score closer to -1 represented down-regulation of the pathway. We calculated GSVA scores for all pathways present in Reactome^14^, KEGG^15^ and biocarta^16^ pathways. We then tested whether there is a difference between the GSVA enrichment scores from different pair of phenotypes using a linear model and moderated t-statistic.

As examples, we found that 605, 589, and 529 (**Supplementary Table 2a-c**) pathways were up-regulated in IDH-wt glioblastomas, IDH-mut astrocytomas and oligodendrogliomas respectively compared to healthy brain samples from GTEx. A total of 456 pathways (**Supplementary Table 2d**) were up regulated in all three adult glioma subtypes compared to healthy brain. The top pathways which were enriched in all adult glioma subtypes were pathways enriched for cell cycle, DNA repair, translation, splicing, oncogenic signaling pathways such as RAS pathway, Notch pathway, MHC pathway, PI3K/AKT Signaling, Wnt pathway, SHH pathway (**Fig. 5, Supplementary Fig. 5a**). Additionally, neurotransmitter pathways, calcium channel, potassium channel opening pathways were up regulated in healthy brain regions compared to pediatric tumors and adult gliomas (**Supplementary Fig. 5b**).

**Fig 5.**
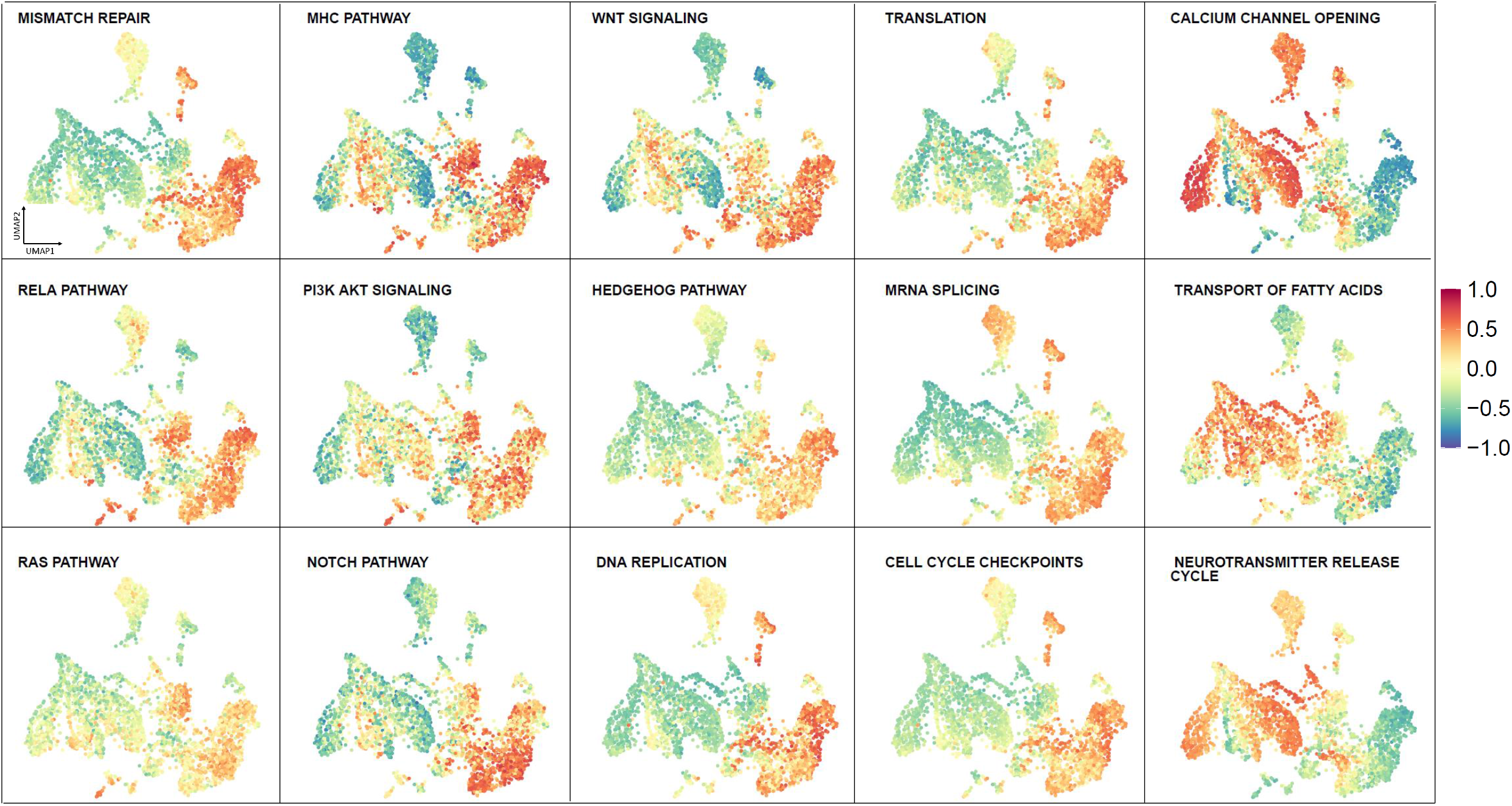
Visualization of GSVA Pathway scores across Brain-UMAP for cancer pathways and cellular processes.

Interestingly, we noted that the two small clusters (IDH mutated grade 2 and grade3 oligodendroglioma and grade 4 IDH-wt glioblastomas) from the CGGA dataset were enriched in pathways related to olfaction, glucoronidation, ascorbate and aldarate metabolism and xenobiotics (**Supplementary Fig. 5c**) in comparison to the main adult glioma cluster.

While visualizing pathways across the reference Brain-UMAP is extremely informative, researchers exploring targets for drug development may be also interested in investigating individual genes of a particular pathway. For example, Reactome’s mismatch repair pathway (R-HAS-5358508) and Biocarta RELA pathway (M10183) were both found to be up-regulated in all adult glioma subtypes compared to healthy normal brain, coloring in the gene expression for individual genes over the Brain-UMAP show different gene expression patterns across members of the same pathway. (**Fig. 6, Supplementary Fig. 6**). For example, almost all the genes in the mismatch repair pathway have elevated gene expression levels in medulloblastoma, except RPA3 and POLD4.

**Fig 6.**
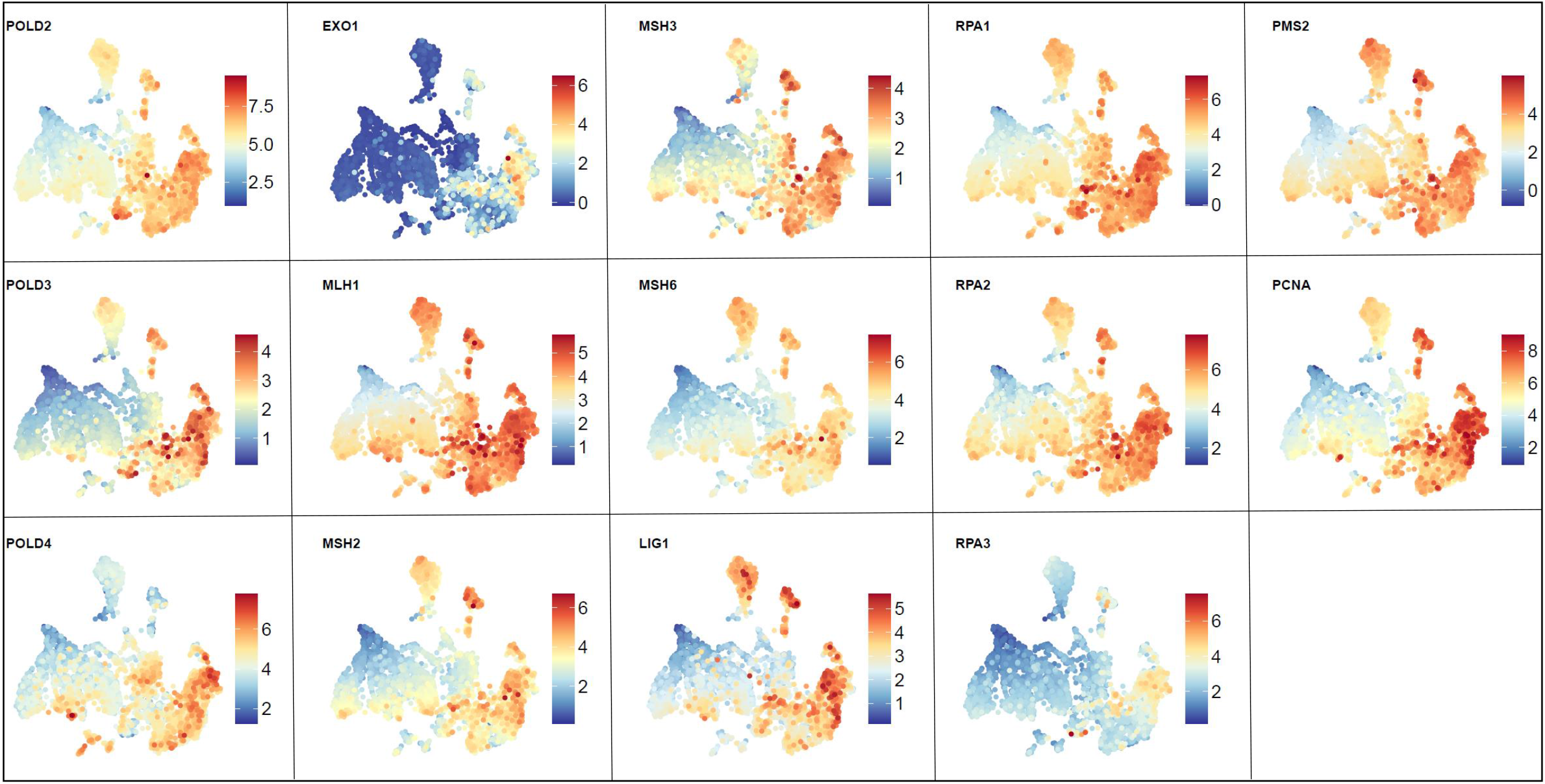
Visualization of gene expression profiles for genes from the Reactome mismatch repair pathway across the Brain-UMAP

### Studying candidate genes at multiple genomic levels

While bulk gene expression exhibited illuminating patterns in our data, analyzing other genomic information such as copy number alteration, gene fusions, or somatic alteration for each of these tumors can further enhance our understanding for a given gene of interest. Since processed copy number, gene fusions and somatic variants were publicly available only for two out of five datasets (TCGA and CBTN), we first built a much smaller UMAP using only the bulk gene expression data from these two datasets. The resulting UMAP (**Fig. 7a**) showed a similar clustering pattern as our original Brain-UMAP. Next, we downloaded the copy number calls, gene fusion calls and somatic variants for adult gliomas from Genomic Data Commons (GDC) and the pediatric tumors from CBTN.

**Fig 7.**
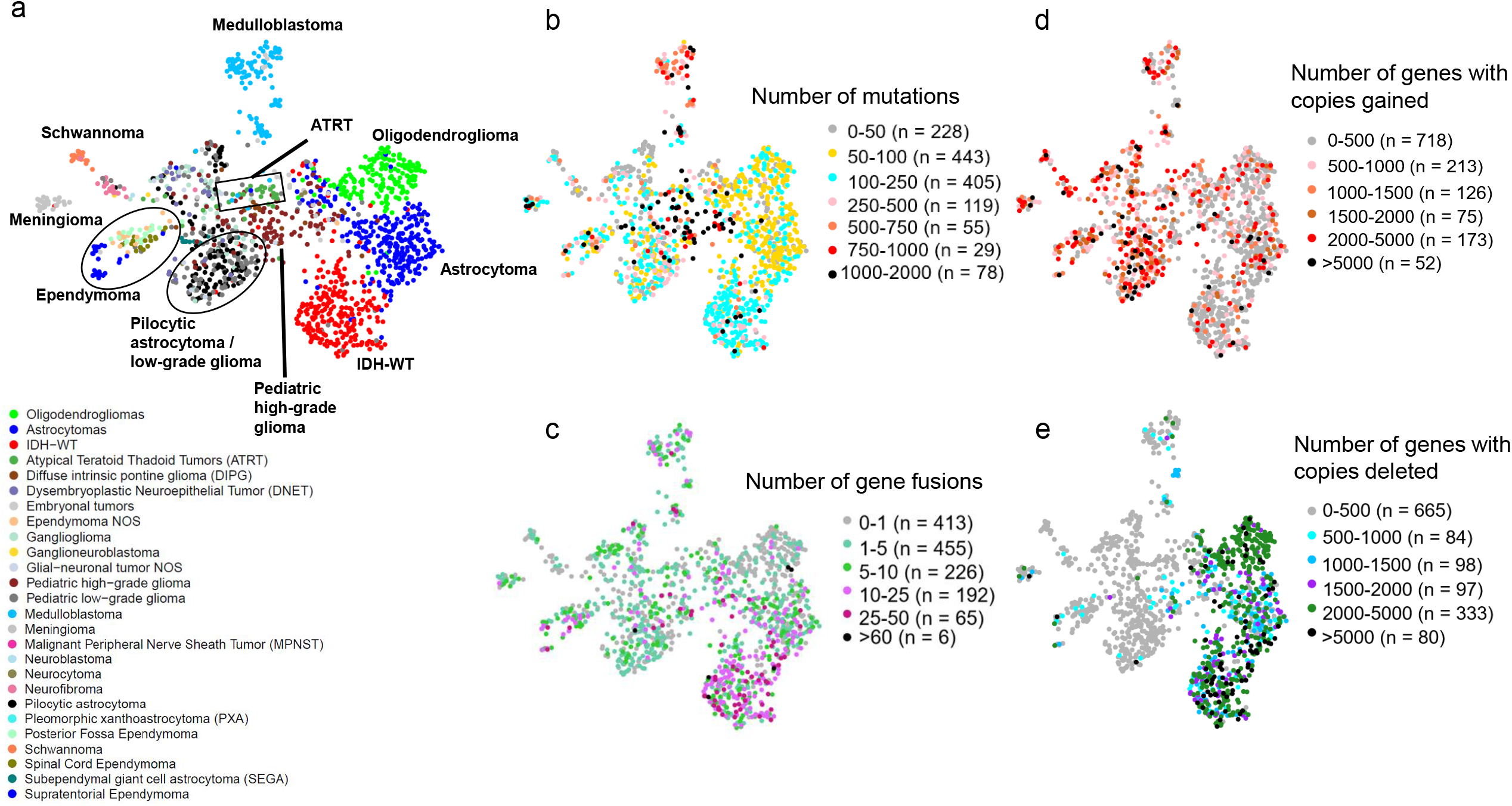
(a) UMAP of pediatric tumors and adult glioma subtypes from TCGA. Coloring in UMAP of pediatric tumors and adult glioma subtypes from TCGA by (b) number of point mutations and (c) number of gene fusions per tumor.

Of note, the IDH-mut oligodendrogliomas, astrocytoma and pediatric high-grade gliomas have the highest mutational burden and number of gene fusions across all brain diseases (**Fig. 7b-c, Supplementary Fig. 7a-b**) in comparison to copy number profiles (**Fig. 7d-e**) which show a mixture of genes with amplified and lost copy number profiles across all brain diseases. Gene fusions are another class of potential oncogenic drivers in cancer, including pediatric cancers.^17^ We observed that specific gene fusions and/or gene fusion partners were enriched in specific cancer subtypes. Adult IDH-wt glioblastoma frequently harbored gene fusions involving EGFR (17.7% of tumors, most commonly EGFR−PSPHP1, EGFR−LINC01445, EGFR−SEC61G−DT), whereas pediatric lower-grade gliomas frequently harbored BRAF fusions (KIAA1549−BRAF in 32% of low-grade gliomas and in 60% of pilocytic astrocytomas). In addition, supratentorial ependymomas most commonly harbored the C11orf95−RELA fusion (71% of tumors, also known as ZFTA-RELA) and meningiomas most frequently harbored fusions in NF2 and YAP1 (both 15% of tumors) **(Supplementary Fig. 7c)**.

Armed with additional genomic information such as gene fusions, copy number variation and somatic alternation, we investigated members of the Reactome mismatch repair pathway (R-HAS-5358508) (**Supplementary Fig. 8a-c**). We observe that different genes belonging to this pathway exhibit different trends, for example genes such as POLD1(chr19), RPA2(chr1), and LIG1(chr19) loose a copy in IDH-mut oligodendrogliomas, while genes RPA3(chr19), PMS2(chr19), PCNA(chr20) and POLD2(chr7) gain a copy in IDH-wt GBM. While most genes show elevated gene expression levels for all brain tumors, of interest is EXO1 and RPA3 which are only up-regulated in IDH-wt GBM samples. While members of this pathway do not form gene fusions, they get mutated in different brain tumors. For example, in pediatric high grade tumors, we observed mutations in all the members of the mismatch repair pathway - MSH2 (22%), MSH6 (15.71%), POLD3 (14.29%), MSH3 (12.86%), LIG1(11.43%), EXO1(10%), PMS2(10%), PCNA(10%), POLD1 (8.57%), RPA2(8.57%), MLH1 (7.14%), RPA1(7.14%), POLD2(5.71%), POLD4(5.71%) and RPA3(5.71%).

Oncogenes show altered gene expression in tumor samples, leading to abnormal phenotype in samples. Understanding gene expression patterns across various cancers of the nervous system can further our understanding of the disease. When studying known oncogenes such as EGFR, PTEN and CIC at a gene level (**Fig. 8**), as expected, we observe high number of mutations, high number of gene fusions, amplified gene expression values and copy number gains for EGFR across IDH-wt GBM. This contrasts with PTEN which shows loss of 1 copy in IDH-wt GBM samples. CIC, a transcriptional repressor, shows high number of mutations and copy number loss in oligodendrogliomas. For pediatric tumors, we observe BRAF gene fusions (**Fig. 8**) in 63% pilocytic astrocytoma tumors and 34% of low grade pediatric tumors (**Supplementary Table 3a**) ALK (**Fig. 8**) mutations are also observed in 38% high-grade pediatric tumors, 33% spinal cord ependymomas, 22% ATRT and 21% of the medulloblastoma tumors (**Supplementary Table 3b**).This reference landscape is a useful research tool for the scientific community, where researchers can explore existing data to increase their understanding of oncogenic pathways and individual genes that make up these pathways, potentially uncovering candidates for novel therapeutic targets. By providing access to this reference landscape via an open source website like Oncoscape, we provide an interactive tool to researchers doing CNS research.

**Fig 8.**
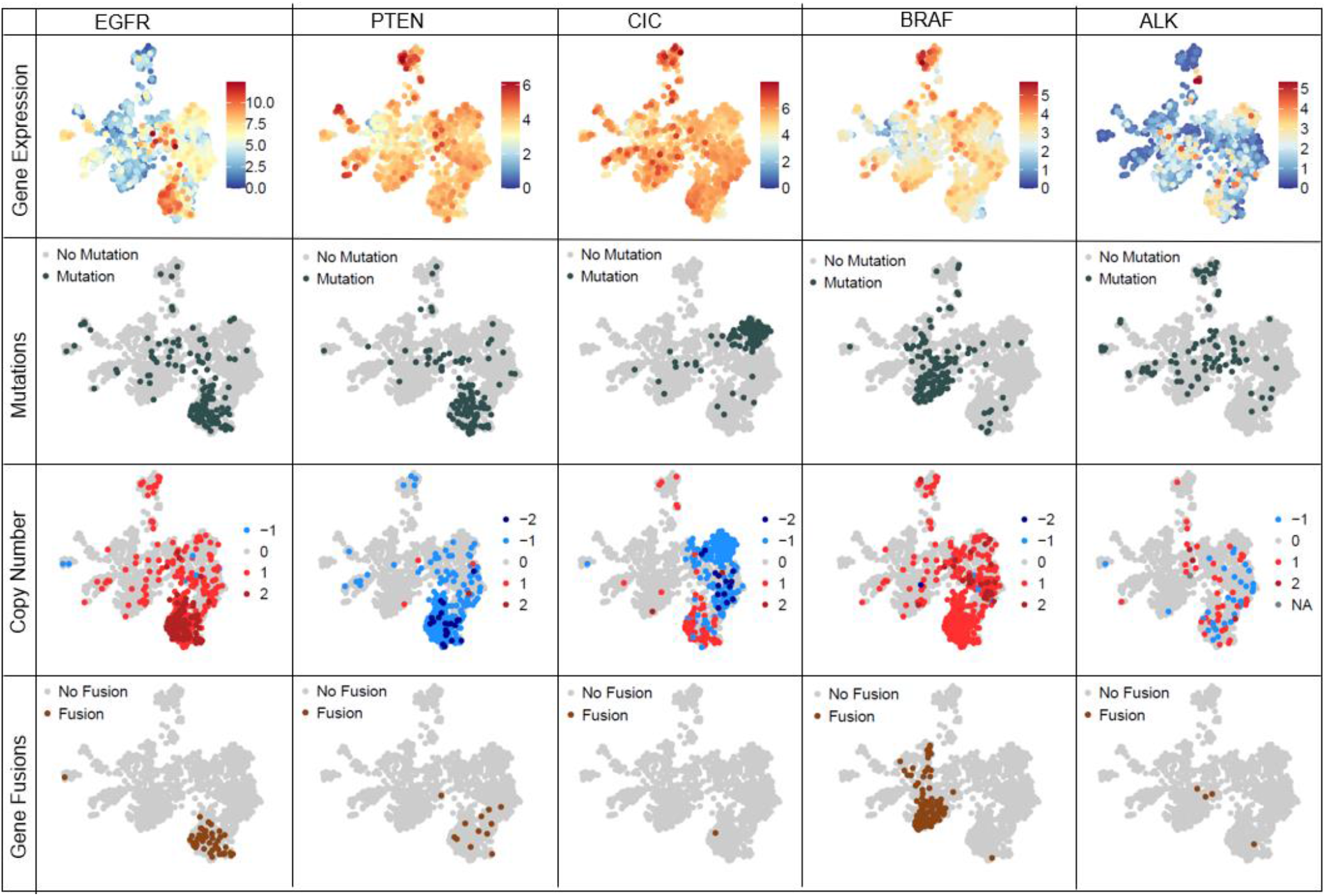
Integration and visualization of genomic information such as gene expression, mutation, copy number and gene fusions at a single gene level across Brain-UMAP for 5 genes – EGFR, PTEN, CIC, BRAF and ALK.

## Discussion

As costs for performing RNASeq continues to decline with increasing technological advances, more and more tumors will be sequenced, and additional tumor banks will be created with the underlying goal to understanding cancer’s complexity. The wealth of knowledge that that already exists in publicly available datasets such as those described here (GTEx, TCGA, CGGA, CBTN) is remarkable. Individually these datasets comprise of well-defined biologically similar set of patient samples and allow the analysis in exquisite resolution of genetic changes from one sample to the next. For example, comparing medulloblastomas to other medulloblastomas allows for precise characterization of molecular subtypes. As researchers, we can get so focused on comparing like with like that we lose sight of the proverbial forest for focusing too much on the leaves of a single tree. By integrating multiple datasets while correcting for batch effects, such as with the Brain-UMAP presented here, we can harness the power of multiple datasets.

This landscape can be used to prospectively compare patients, similar to work that has been done using methylation arrays. Unlike methylation arrays, however, RNA seq is based on gene expression and therefore each cluster represented on the landscape contains granular information about the underlying tumor biology of the samples it contains. Because the overall expression pattern is identifiable for each tumor, this allows for cross comparison of various kinds of cancer to allow for characterization of tumor types not previously known. For example, asking questions about expression of particular genes or pathways across a wide panel of samples and tumor cohorts may uncover previously unknown roles or similarities between adult and pediatric tumor subtypes, which can possibly open new avenues for therapeutic discovery.

The landscape that we present reveals correlations between tumor type and outcome. We have learned how diagnosis of a given tumor type can be confirmed using various pieces of genomic information. For example – the adult glioma subtype of oligodendroglioma can be confirmed by presence of co-deletion of chr1p/19q and somatic alteration in IDH1.

We are also able to compare tumor samples which were sequenced years apart (TCGA and CGGA) in two different continents and confirm that these differences did not contribute to expression changes in majority of the tumor samples. We are also able to identify novel entities and gain insight into genes that drive their unique character, such as in the case of the two adult glioma subtypes seen only in the CCGA data that appear to strongly and uniquely express genes involved in olfaction.

Using this study as an example, we have seen how one diagnosis can be comprised of more than one expression entity, such as in the case of medulloblastoma and ependymoma. The UMAP also indicates room for better classification of tumors. For example, we found tumors which were documented as one type in the publicly available database, but in our UMAP were found to cluster with tumors of a different type. For example, some embryonal tumors ended up clustering with the medulloblastoma samples indicating that they may be medulloblastomas, or at least share many common features with medulloblastoma. We also have seen that multiple diagnoses may really be one entity as in the case of tumors diagnosed as either pilocytic astrocytoma (PA) or pediatric low-grade glioma.

Reference landscapes like the Brain-UMAP, will be informative for researchers who wish to obtain a quick diagnosis or characterization of newly obtained tumor samples. For example, if we look at the medulloblastoma subtypes from CBTN, there are 14 group 3, 49 group 4, 30 SHH, 9 WNT and 19 unclassified medulloblastoma samples. For the 19 unclassified samples, we can predict which subtype they belong to, based on which medulloblastoma subtype samples they cluster with. (Supp Fig 4a).

By combining results from both RNA-seq data (gene expression, gene fusions) as well as whole genome sequencing (copy number, mutation calls) and with the help of a few examples, we illustrate the knowledge that can be mined from this resource, at both a pathway level as well as a gene level. The work presented here demonstrates the utility of reference landscapes for combining genomic data across multiple tumor types for both diagnosis, prognosis and better understanding the biology of the tumors that are similar to a given patient collected prospectively. The power of visualizing gene expression changes, regulation of pathways, chromosomal alteration, gene fusions across multiple tumor types is informative to every researcher, especially those who may not have immediate access to computational biology experts. The reference landscape described here provides a useful tool for researchers interested in gene level questions across large scale patient data, while the methods used for integrating data sources highlight the tremendous potential for combining future datasets with existing resources to address complex biological questions.

## Methods

All analyses were performed in R (https://www.r-project.org/) using Bioconductor (https://www.bioconductor.org/) packages. We have deposited all scripts, associated data, at: https://zenodo.org/badge/latestdoi/584982012 to maximize transparency and reproducibility. TCGA data from GDC was downloaded using R/Bioconductor package TCGAbiolinks^18,19^, SummarizedExperiment^20^ was used to store adult glioma and pediatric tumor data. All plots were made using ggplot2^21^, RcolorBrewer^22^.

### Obtaining gene expression RNASeq Data

RNA Seq gene expression was downloaded from two sources for adult gliomas, GTEx defined healthy brain samples and pediatric tumors respectively (Supplemental Table 1).

### Conversion of abundance estimates to transcripts per million (TPM)

For consistency, we converted all FPKM gene expression data to TPM data using the formula

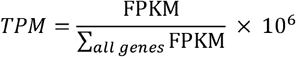

as described by Collins et al^23^.

### Uniform Manifold Approximation and Project (UMAP)

Uniform Manifold Approximation and Project (UMAP) values were generated using the R function umap() using all protein-coding genes and visualized with R package ggplot2^21^.

### Obtaining Copy Number

Gistic2 thresholded gene level copy number variation estimated using the GSITIC2 method were downloaded from UCSC Xena(https://xenabrowser.net/datapages/?cohort=TCGA%20Lower%20Gra_de%20Glioma%20(LGG)&removeHub=https%3A%2F%2Fxena.treehouse.gi.ucsc.edu%3A443 and https://xenabrowser.net/datapages/?cohort=TCGA%20Glioblastoma%20(G_BM)&removeHub=https%3A%2F%2Fxena.treehouse.gi.ucsc.edu%3A443) for adult gliomas. Copy Number calls for pediatric tumors estimated from the GISTIC2 pipeline were downloaded from https://github.com/AlexsLemonade/OpenPBTA-analysis.

### Obtaining Gene Fusions

Gene fusions for adult gliomas estimated using Arriba and Star-Fusion as per the GDC RNA-Seq pipeline were downloaded from the GDC portal (https://portal.gdc.cancer.gov/projects/TCGA-GBM, https://portal.gdc.cancer.gov/projects/TCGA-LGG). Gene fusions for pediatric tumors were downloaded from https://github.com/AlexsLemonade/OpenPBTA-analysis. Only high-confidence gene fusion calls were retained from ARRIBA.

### Obtaining Mutations

Annotated Variant call Format (VCF) files containing somatic variants from MuSE, VarScan2, MuTect2 and Somatic Sniper were downloaded from GDC portal for TCGA-GBM and TCGA-LGG. The variants for each patient were combined based on custom script present in our github repository. Somatic Variant calls for pediatric tumors was obtained from https://github.com/AlexsLemonade/OpenPBTA-analysis.

### GSVA Pathway Analysis

Gene sets for all the pathways from Biocarta, Kyoto Encylcopedia of Genes and Genomes (KEGG) and Reactome pathways were downloaded from Molecular Signature Databases (MSigDB)(v7.2)^24^. https://www.gsea-msigdb.org/gsea/msigdb/collections.jsp#C2 Batch corrected log2(TPM) counts from each pipeline were used to conduct a Gene Set Variation Analysis (GSVA)^25^ using Biocarta, KEGG and Reactome pathways. GSVA scores obtained from 1 and -1 for each sample, were visualized using ggplot2.

### Kaplan-Meier curves for TCGA and CGGA patients

To generate Kaplan-Meier curves for TCGA and CGGA patients, all samples labeled recurrent, secondary, or normal tissue were removed. Duplicate samples were also excluded so that each patient was represented by exactly one sample. Kaplan-Meier curves were drawn by the Python package *lifelines* (Python version 3.8.6, *lifelines* version 0.25.2)^26^ using survival data and glioma subtype labels downloaded from GDC, where the world health organization (WHO) 2016 criteria for the classification of adult diffuse gliomas was used to determine glioma subtype^27^.

### Survival-Annotated UMAP

To annotate the Brain-UMAP with survival data, each TCGA and CGGA sample was colored with the median survival of a cohort of patients (nearest neighbors) close to the sample in question on the UMAP landscape. The notion of nearest neighbors was defined as a number of samples within a radius of 2 from the sample in question under the constraint that all nearest neighbors must be of the same glioma subtype and from the same dataset (TCGA or CGGA) as the sample in question. The radius parameter was chosen qualitatively. The number of nearest neighbors was defined as 25% of the total number of samples of the same glioma subtype and dataset as the sample in question. If the median survival of a sample’s cohort of nearest neighbors is undefined, or if a sample has fewer than 10 nearest neighbors, the point is not colored in. Non-primary and duplicate samples were excluded in the same manner as was done for the Kaplan-Meier curve analysis.

### Oncoscape integration

Gene expression and survival data analysis result files were prepared according to the oncoscape instructions here: https://github.com/FredHutch/OncoscapeV3/blob/master/docs/upload.md

### Data Analysis

All statistical analyses and plots were done in R (v.3.3.1) as implemented in Rstudio (v.1.0.136). Plots were created using the R basic graphics. The following R packages were used: GenomicAlignments53 (v.1.24), reshape64 (v.0.8.8), png65 (v.0.1–7), ape66 (v.5.3) and seqinr67 (v.3.6–1).

## Acknowledgements

We thank members of the Holland Lab at Fred Hutchinson Cancer Research Center for discussions. This research was supported by the R35 CA253119-01A1(E.C.H.), NIH U54 CA193461 (E.C.H.), National Institutes of Health R01 CA195718 (E.C.H.), R01 CA100688 (E.C.H.), T32 CA9657-25 (S.S.P.), U54 DK106829 (S.S.P.), R21 CA223531 (S.S.P.); K22 CA258953-01 (SSP), Jacobs Foundation Research Fellowship (S.S.P.) and National Science Foundation Graduate Research Fellowship Program DGE-1762114 (N.N.).

## Author Information

### Contributions

S.A., S.S.P., F.S., and E.C.H. conceived and organized the study based on S.A.’s proposed design. S.A. and N.N. performed all the analysis. M.J. incorporated the data into oncoscape. S.A. and S.S.P. wrote the manuscript, with contributions from all authors.

Corresponding Author

Correspondence to Eric C Holland

### Data availability

All data are available in the main text, supplementary materials and/or accompanying database is found at https://zenodo.org/badge/latestdoi/584982012. We used data from recount2 (https://jhubiostatistics.shinyapps.io/recount/), CBTN (https://github.com/AlexsLemonade/OpenPBTA-analysis), copy number data for TCGA-GBM (https://xenabrowser.net/datapages/?cohort=TCGA%20Glioblastoma%20(G_BM)&removeHub=https%3A%2F%2Fxena.treehouse.gi.ucsc.edu%3A443) and TCGA-

LGG(https://xenabrowser.net/datapages/?cohort=TCGA%20Lower%20Gra_de%20Glioma%20(LGG)&removeHub=https%3A%2F%2Fxena.treehouse.gi.ucsc.edu%3A443), Gene fusions from GDC Data portal (https://portal.gdc.cancer.gov/projects/TCGA-GBM, https://portal.gdc.cancer.gov/projects/TCGA-LGG)

### Code availability

All custom code used in this study is available at https://zenodo.org/badge/latestdoi/584982012

## References

1. Cancer Genome Atlas Research, N., et al. The Cancer Genome Atlas Pan-Cancer analysis project. Nat Genet 45, 1113–1120 (2013).

2. Zhao, Z., et al. Chinese Glioma Genome Atlas (CGGA): A Comprehensive Resource with Functional Genomic Data from Chinese Glioma Patients. Genomics Proteomics Bioinformatics 19, 1–12 (2021).

3. Ijaz, H., et al. Pediatric high-grade glioma resources from the Children’s Brain Tumor Tissue Consortium. Neuro Oncol 22, 163–165 (2020).

4. Carithers, L.J., et al. A Novel Approach to High-Quality Postmortem Tissue Procurement: The GTEx Project. Biopreserv Biobank 13, 311–319 (2015).

5. McFerrin, L.G., et al. Analysis and visualization of linked molecular and clinical cancer data by using Oncoscape. Nat Genet 50, 1203–1204 (2018).

6. Shapiro, J.A., et al. OpenPBTA: An Open Pediatric Brain Tumor Atlas. bioRxiv (2022).

7. Collado-Torres, L., et al. Reproducible RNA-seq analysis using recount2. Nat Biotechnol 35, 319–321 (2017).

8. Subramanian S A.T. Childhood Brain Tumors. In: StatPearls [Internet]. Treasure Island (FL): StatPearls Publishing, https://www.ncbi.nlm.nih.gov/books/NBK535415/ (2022).

9. Arora, S., Pattwell, S.S., Holland, E.C. & Bolouri, H. Variability in estimated gene expression among commonly used RNA-seq pipelines. Sci Rep 10, 2734 (2020).

10. Johnson, W.E., Li, C. & Rabinovic, A. Adjusting batch effects in microarray expression data using empirical Bayes methods. Biostatistics 8, 118–127 (2007).

11. Pattwell, S.S., et al. A kinase-deficient NTRK2 splice variant predominates in glioma and amplifies several oncogenic signaling pathways. Nat Commun 11, 2977 (2020).

12. Bolouri, H., Zhao, L.P. & Holland, E.C. Big data visualization identifies the multidimensional molecular landscape of human gliomas. Proc Natl Acad Sci U S A 113, 5394–5399 (2016).

13. Pollack, I.F., Agnihotri, S. & Broniscer, A. Childhood brain tumors: current management, biological insights, and future directions. J Neurosurg Pediatr 23, 261–273 (2019).

14. Gillespie, M., et al. The reactome pathway knowledgebase 2022. Nucleic Acids Res 50, D687–d692 (2022).

15. Kanehisa, M. & Goto, S. KEGG: kyoto encyclopedia of genes and genomes. Nucleic Acids Res 28, 27–30 (2000).

16. BioCarta. Biotech Software & Internet Report 2, 117–120 (2001).

17. Szulzewsky, F., et al. Both YAP1-MAML2 and constitutively active YAP1 drive the formation of tumors that resemble NF2 mutant meningiomas in mice. Genes Dev 36, 857–870 (2022).

18. Colaprico, A., et al. TCGAbiolinks: an R/Bioconductor package for integrative analysis of TCGA data. Nucleic Acids Res 44, e71 (2016).

19. Silva, T.C., et al. TCGA Workflow: Analyze cancer genomics and epigenomics data using Bioconductor packages. F1000Res 5, 1542 (2016).

20. Morgan M O.V., Hester J, Pagès H. SummarizedExperiment: SummarizedExperiment container. R package version 1.16.0. (2019).

21. Wickham, H. ggplot2: Elegant Graphics for Data Analysis, (Springer-Verlag New York, 2016).

22. Neuwirth, E. Package ‘RColorBrewer’, ColorBrewer Palettes. (2014).

23. Bo Li, C.N.D. RSEM: accurate transcript quantification from RNA-Seq data with or without a reference genome. BMC Bioinformatics (2011).

24. Aravind Subramanian, P.T., Vamsi K. Mootha, Sayan Mukherjee, Benjamin L. Ebert, Michael A. Gillette, Amanda Paulovich, Scott L. Pomeroy, Todd R. Golub, Eric S. Lander, Jill P. Mesirov. Gene set enrichment analysis: A knowledge-based approach for interpreting genome-wide expression profiles. PNAS (2005).

25. Sonja Hänzelmann, R.C.J.G. GSVA: gene set variation analysis for microarray and RNA-Seq data. BMC Bioinformatics (2013).

26. Davidson-Pilon, C. lifelines: survival analysis in Python. Journal of Open Source Software 4(2019).

27. Louis, D.N., et al. The 2016 World Health Organization Classification of Tumors of the Central Nervous System: a summary. Acta Neuropathol 131, 803–820 (2016).

