## Supplemental Figures for "An RNA seq-based reference landscape of human normal and neoplastic brain"

#### Supplementary Figures

**Supplementary Figure 1.** (a) Principle Component Analysis ( PCA), t-distributed stochastic neighbor embedding (t-SNE) , and Uniform Manifold Approximation and projection for dimension reduction (UMAP) on non-batch corrected and batch corrected datasets from GTEx, TCGA-LGG, TCGA-GBM , CGGA and CBTN. (b) UMAP projection for RNA-seq data from GTEx colored in by postmortem interval (PMI), age (years), sex, Hardy score , type of sample preparation and RNA integrity numbers (RIN).

**Supplementary Figure 2.** UMAP projection for RNA-seq data from TCGA-LGG and TCGA-GBM colored in by (a) gender (b) gain of chromosome 19 and 20 (c) supervised DNA methylation subtype (d) transcription subtype (e) MGMT promoter status (f) TERT promoter status.

**Supplementary Figure 3.** Prediction of survival time (in years) using nearest neighbors, using different numbers of nearest neighbors 10, 15, 20, 50, 75 and 100.

**Supplementary Figure 4.** UMAP projection for RNASeq data from GTEx, CBTN, CGGA and TCGA datasets, Only medulloblastoma subtypes group 3 (MG Group3), group4 (MB Group4), Sonic hedgehog (MB SHH), Wnt (MB WNT) and unclassified medulloblastoma (MB unclassified) samples are colored in. All other samples are greyed out.

**Supplementary Figure 5.** Venn diagram represents the number of pathways that were significantly up-regulated in each of the adult glioma subtypes from TCGA compared to GTEx data. UMAP projection for RNASeq data from GTEx, CBTN, CGGA and TCGA datasets colored in for different pathways (a) up-regulated in adult glioma subtypes, (b) up-regulated in normal brain GTEx samples and (c) up-regulated in the two small clusters (IDH mutated grade 2 and grade3 oligodendroglioma and grade 4 IDH-wt glioblastomas) from the CGGA dataset.

**Supplementary Figure 6.** UMAP projection for RNASeq data from GTEx, CBTN, CGGA and TCGA datasets colored in for genes from the BIOCARTA RELA pathway.

**Supplementary Figure 7.** Boxplots for adult glioma subtypes and pediatric tumor types showing frequency of (a) mutations and (b) gene fusions. (c) Barplots showing percentage of tumors containing top 5 gene fusions in TCGA IDH-wild type , Pediatric low-grade gliomas, pilocytic astrocytoma , meningioma and supratentorial ependymoma.

**Supplementary Figure 8 (a-c).** UMAP projection for RNASeq data from only CBTN and TCGA datasets colored in for genes from the REACTOME MISMATCH REPAIR pathway.

**Supplementary Tables**

Supplementary Table 1a. Table showing number of samples in each of the 12 GTEx defined brain regions.

Supplementary Table 1b. Table showing number of samples for each of the included pediatric tumors from CBTN.

Supplementary Table 1c. Overview of different data sources and processing.

Supplementary Table 2a. List of pathways up-regulated in IDH wild-type glioma subtype compared to normal brain.’

Supplementary Table 2b. List of pathways up-regulated in oligodendrogliomas compared to normal brain.

Supplementary Table 2c. List of pathways up-regulated in IDH mutated glioma subtype compared to normal brain.

Supplementary Table 2d. List of pathways up-regulated in all 3 adult glioma subtypes (IDH wild-type , oligodendroglioma, IDH mutated) compared to normal brain.

Supplementary Table 3a : Number of mutation in BRAF for each tumor type

Supplementary Table 3b: Number of mutation in ALK for each tumor type

**Supplementary Fig 1a**

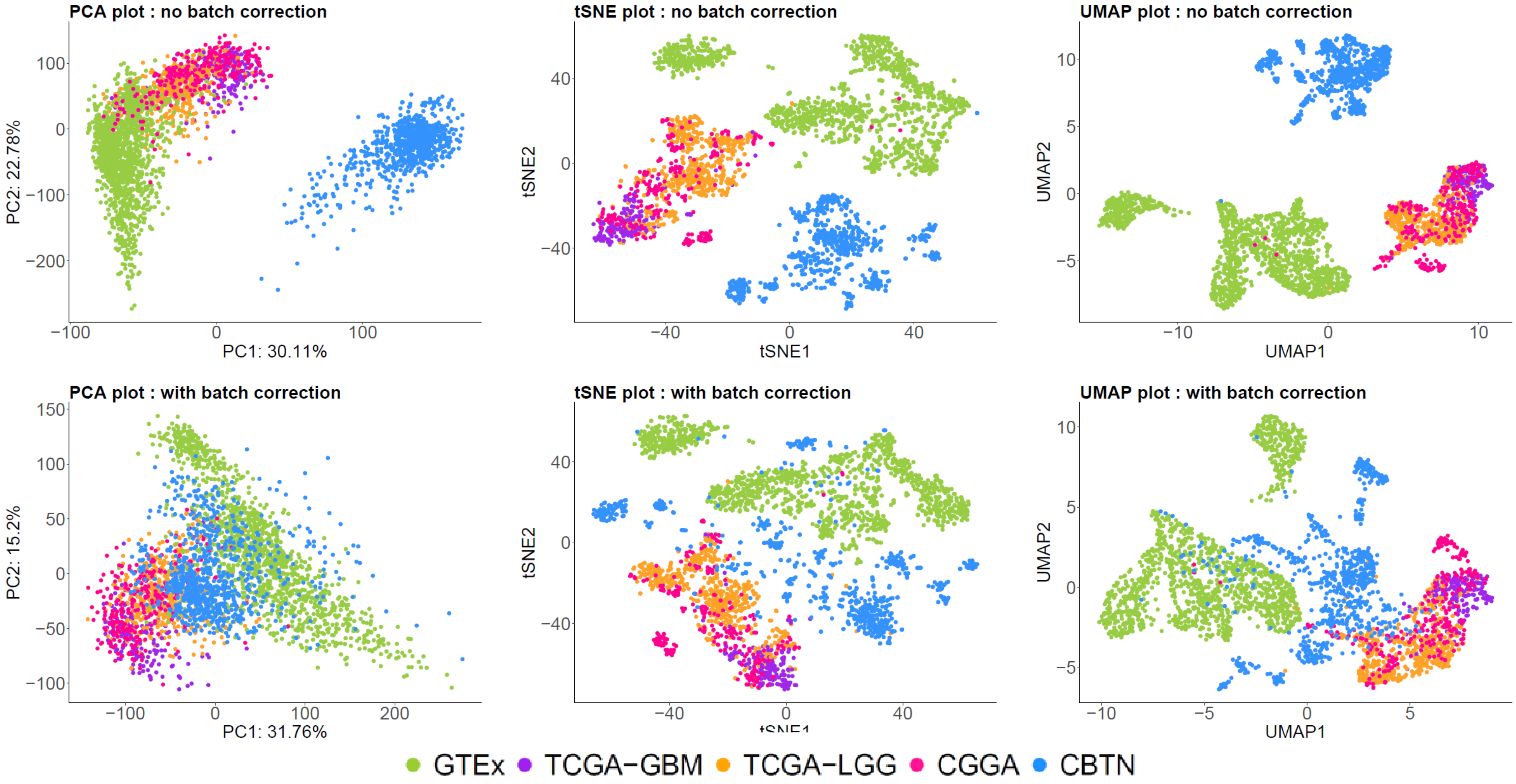


**Supplementary Figure 1.**(a) Principle Component Analysis ( PCA), t-distributed stochastic neighbor embedding (t-SNE) , and Uniform Manifold Approximation and projection for dimension reduction (UMAP) on non-batch corrected and batch corrected datasets from GTEx, TCGA-LGG, TCGA-GBM , CGGA and CBTN.

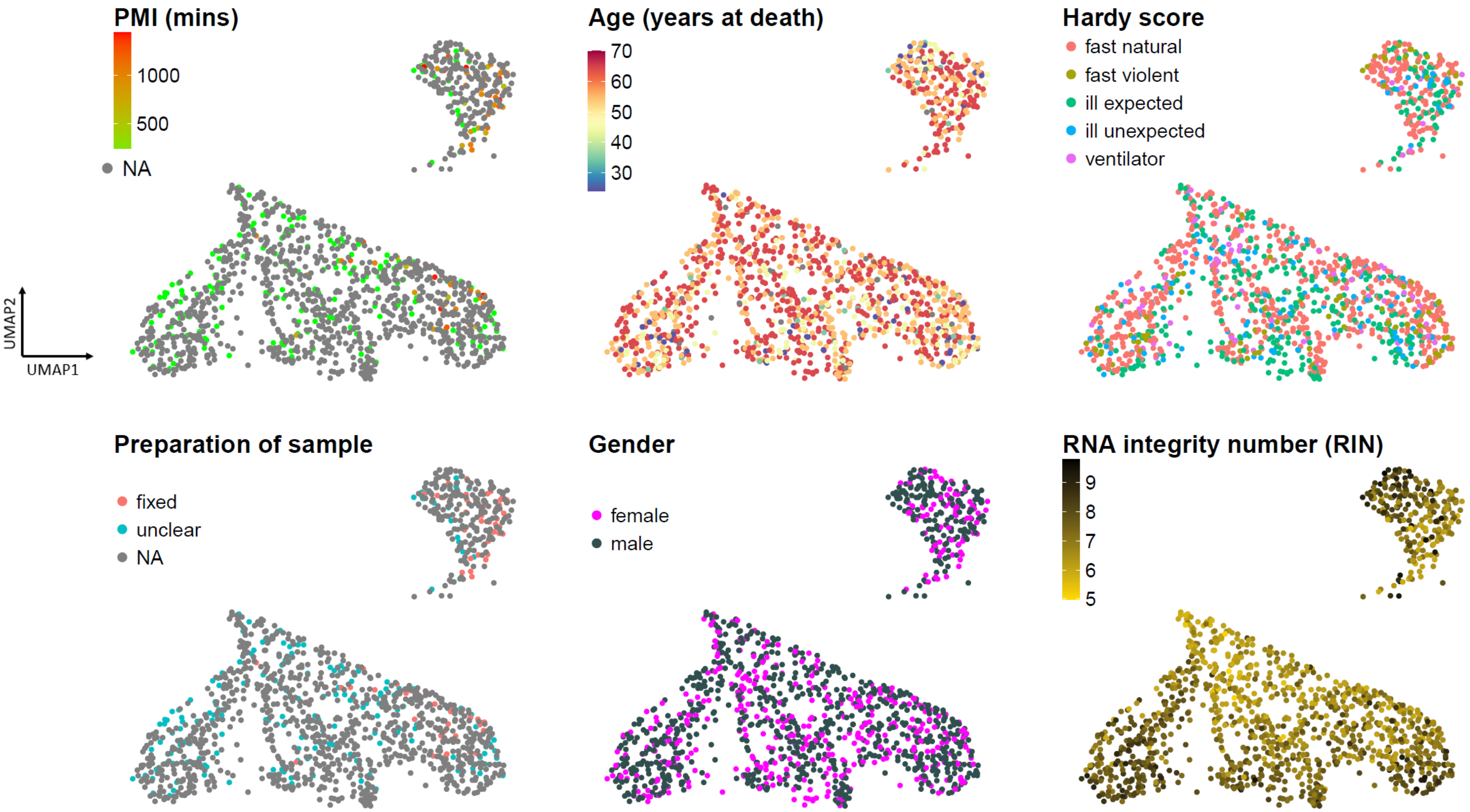


**Supplementary Figure 1.** (b) UMAP projection for RNA-seq data from GTEx colored in by postmortem interval (PMI), age (years), sex, Hardy score , type of sample preparation and RNA integrity numbers (RIN).

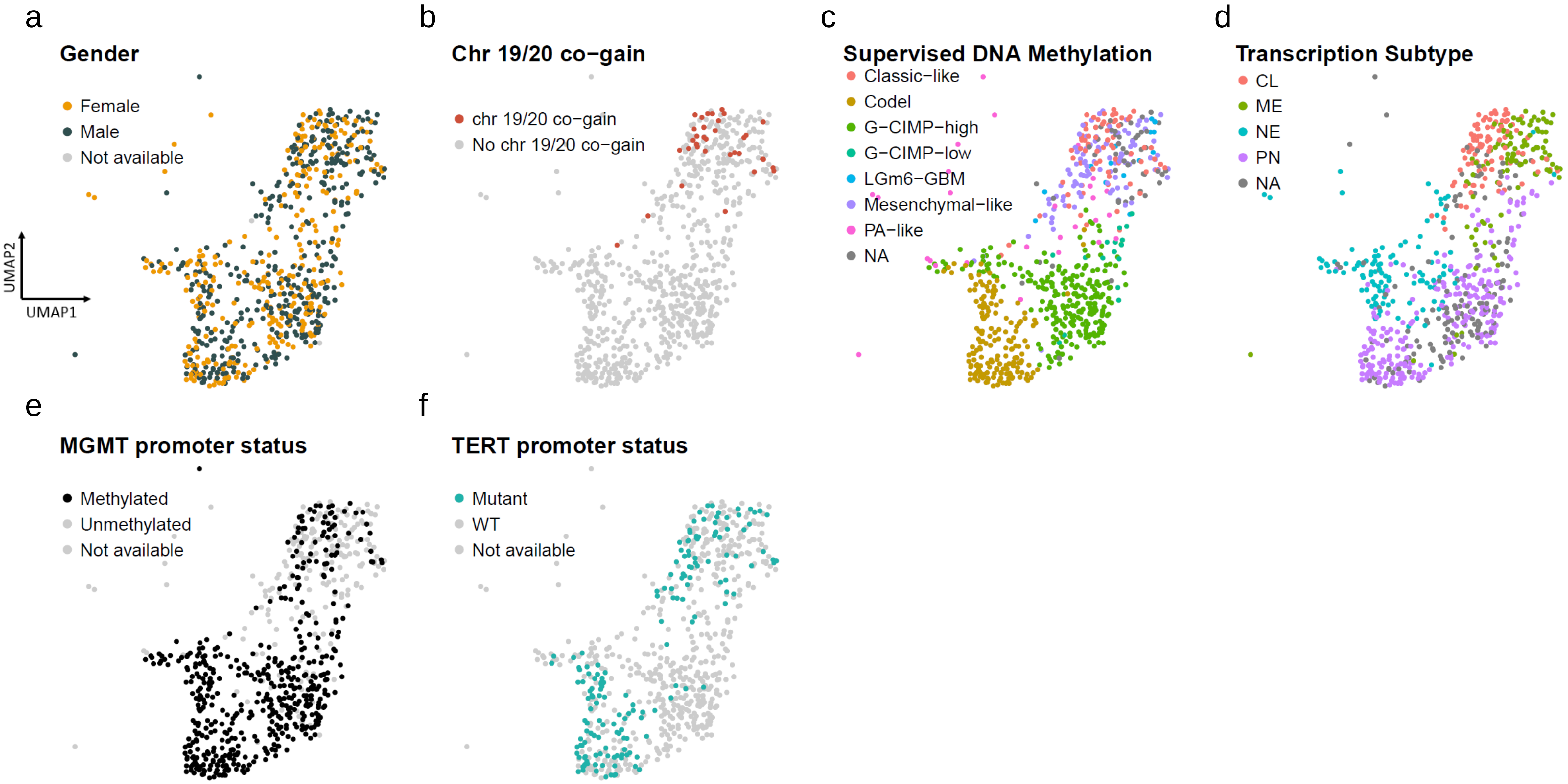


**Supplementary Figure 2.** UMAP projection for RNA-seq data from TCGA-LGG and TCGA-GBM colored in by (a) gender (b) gain of chromosome 19 and 20 (c) supervised DNA methylation subtype (d) transcription subtype (e) MGMT promoter status (f) TERT promoter status.

### Supplementary Fig 3

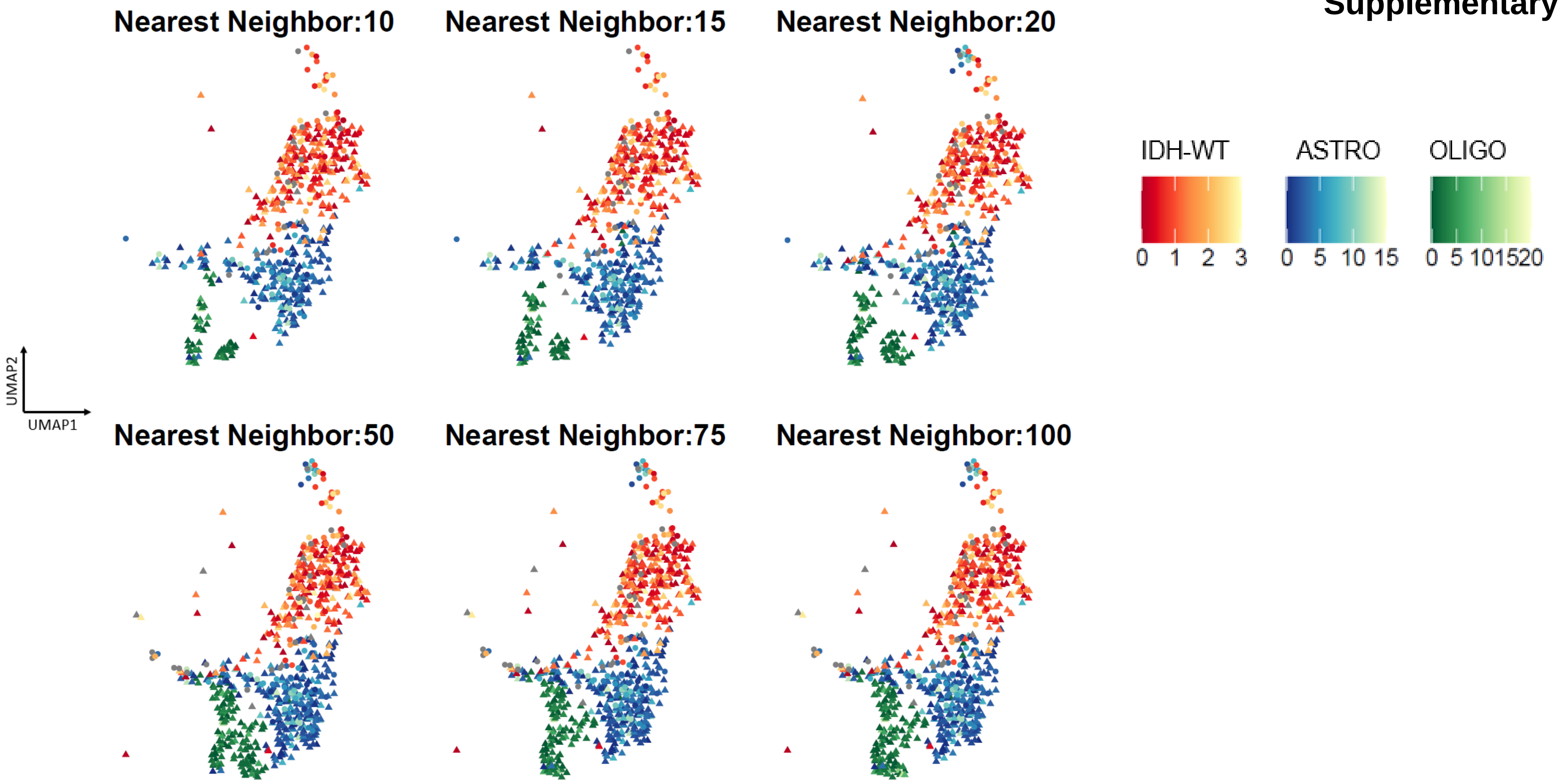


**Supplementary Figure 3.** Prediction of survival time (in years) using nearest neighbors, using different numbers of nearest neighbors 10, 15, 20, 50, 75 and 100.

### Supplementary Fig 4

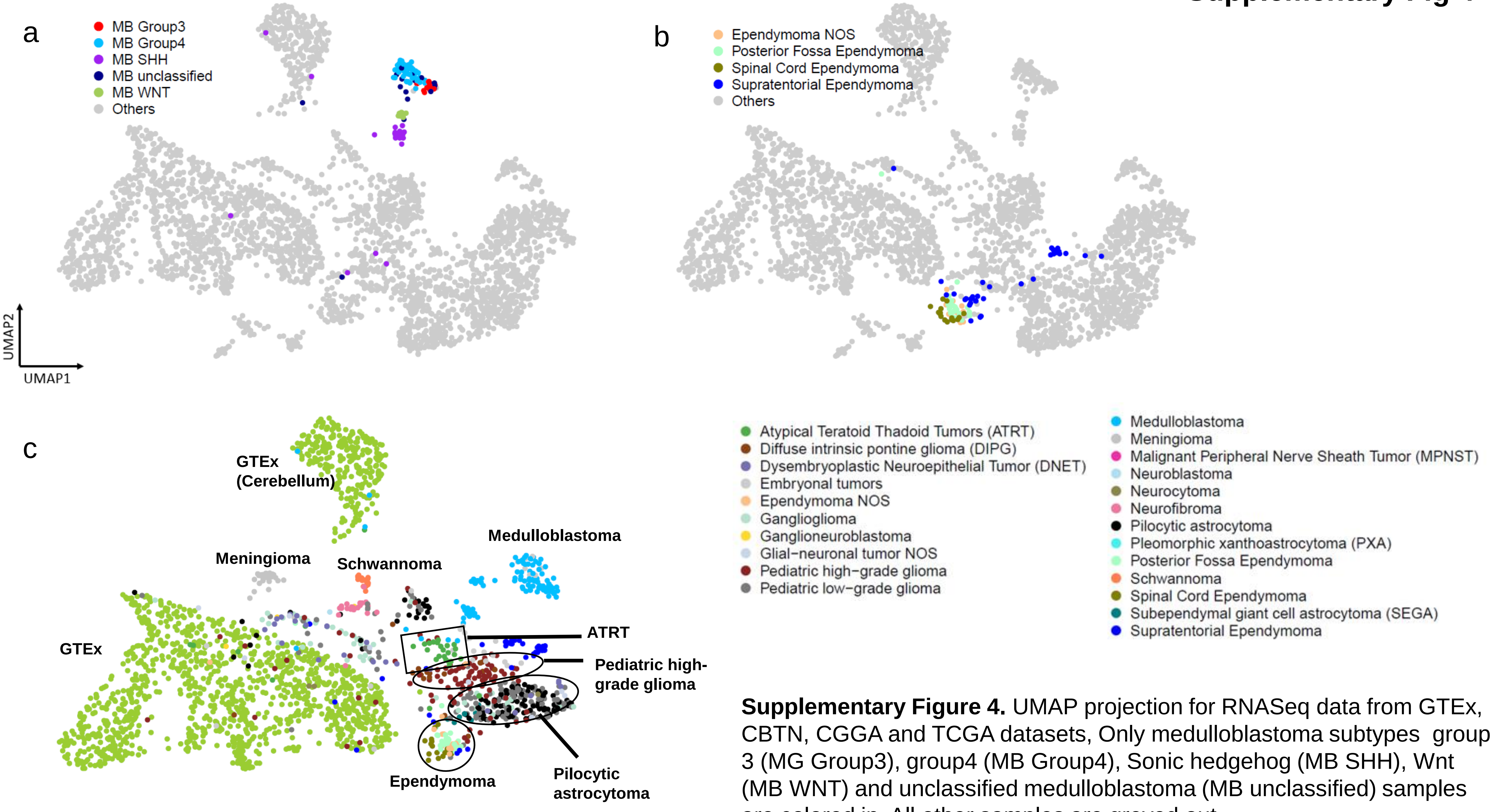


**Supplementary Figure 4.** UMAP projection for RNASeq data from GTEx, CBTN, CGGA and TCGA datasets, Only medulloblastoma subtypes group 3 (MG Group3), group4 (MB Group4), Sonic hedgehog (MB SHH), Wnt (MB WNT) and unclassified medulloblastoma (MB unclassified) samples are colored in. All other samples are greyed out.

**Supplementary Fig 5**

a

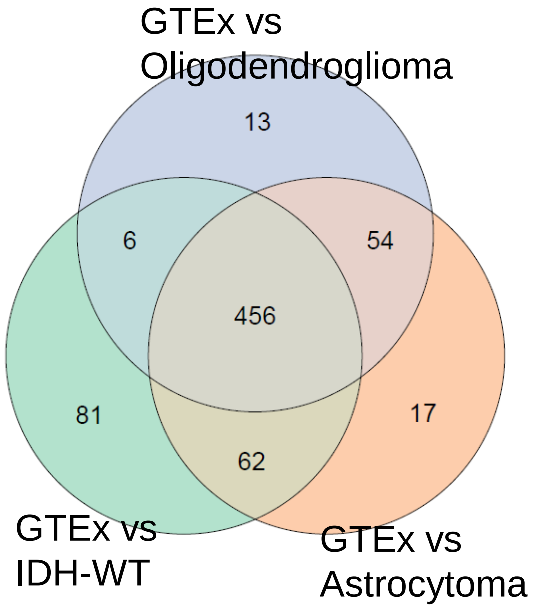


| | Pathway # |
|---|---|
| GTEx vs IDH-WT | 605 |
| GTEx vs Astrocytoma | 589 |
| GTEx vs Oligodendroglioma | 529 |
| GTEx vs all 3 adult glioma subtypes | 456 |

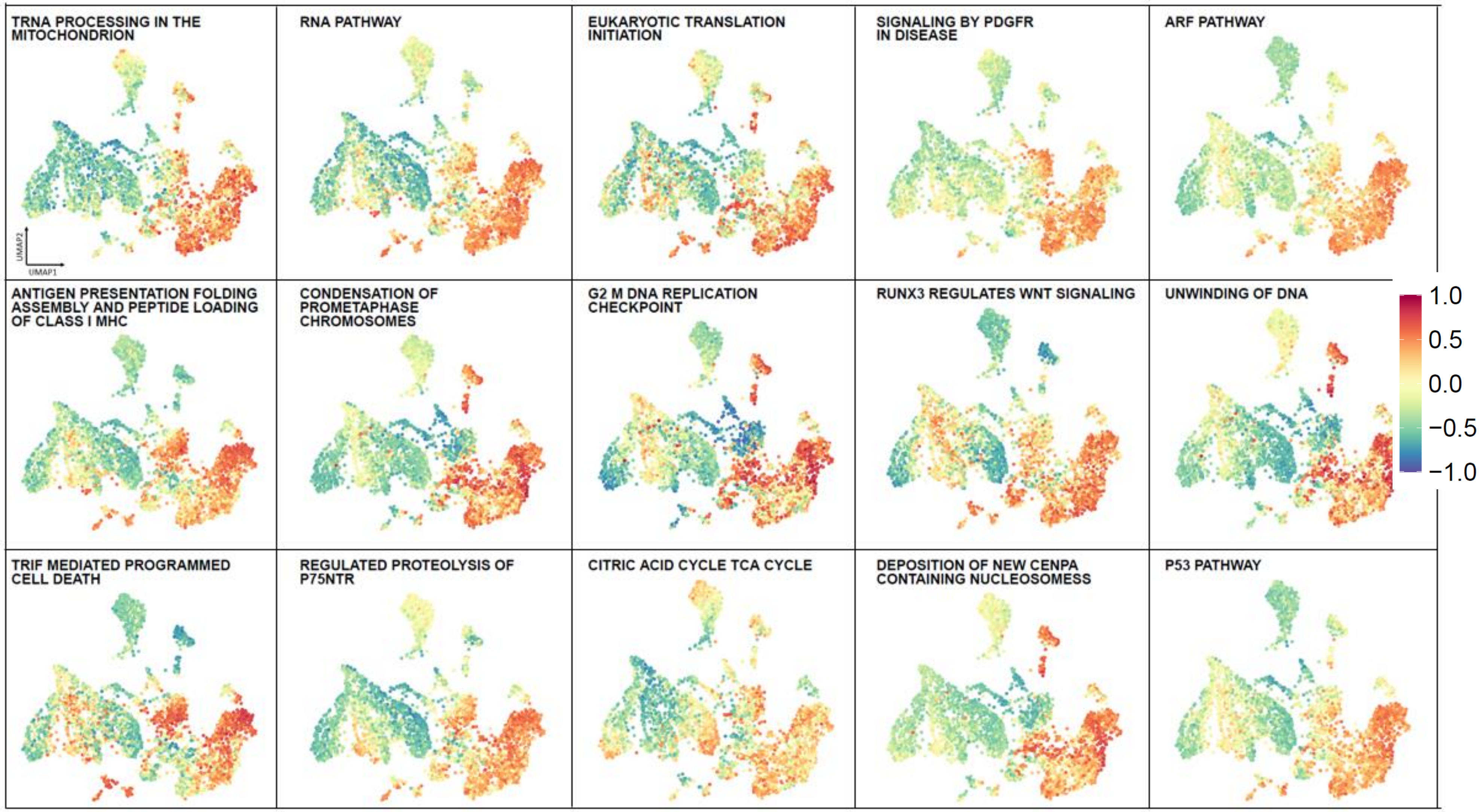


**Supplementary Figure 5.** Venn diagram represents the number of pathways that were significantly up-regulated in each of the adult glioma subtypes from TCGA compared to GTEx data. UMAP projection for RNASeq data from GTEx, CBTN, CGGA and TCGA datasets colored in for different pathways (a) up-regulated in adult glioma subtypes

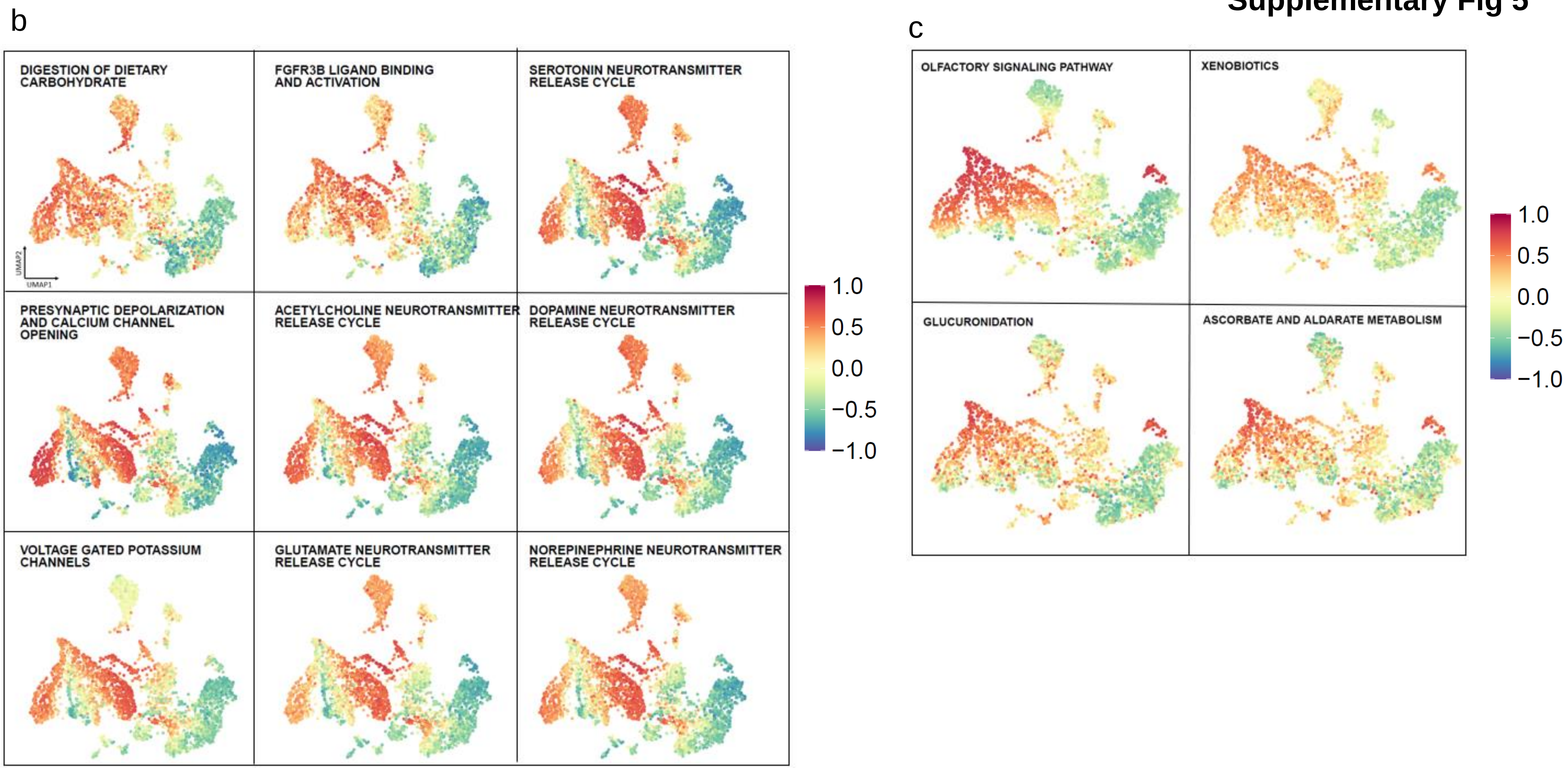


**Supplementary Figure 5.** UMAP projection for RNASeq data from GTEx, CBTN, CGGA and TCGA datasets colored in for different pathways (b) up-regulated in normal brain GTEx samples and (c) up-regulated in the two small clusters (IDH mutated grade 2 and grade3 oligodendroglioma and grade 4 IDH-wt glioblastomas) from the CGGA dataset.

Supplementary Fig 6

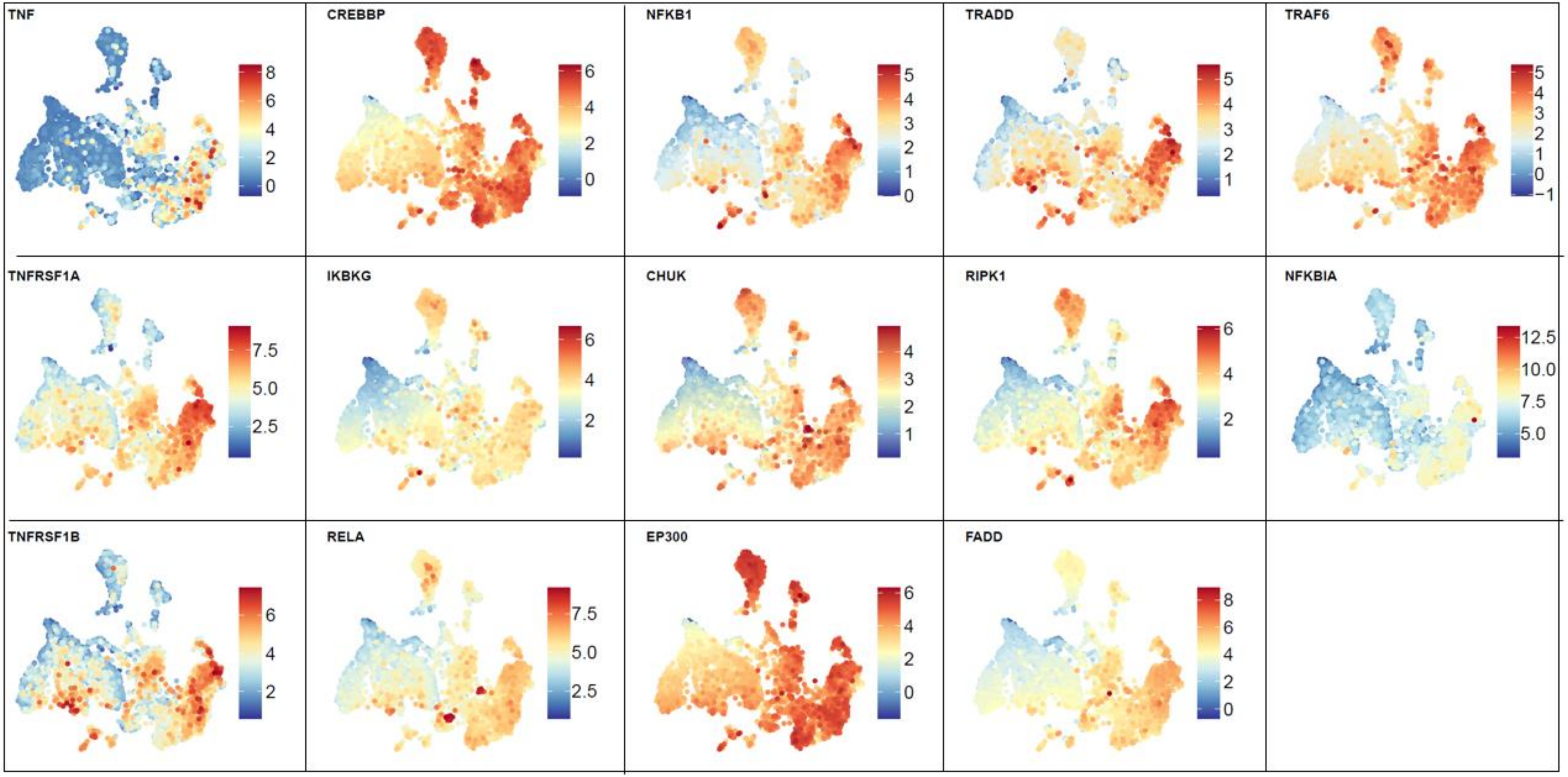


**Supplementary Figure 6.** UMAP projection for RNA-seq data from GTEx, CBTN, CGGA and TCGA datasets colored in for genes from the BIOCARTA RELA pathway.

**Supplementary Fig 7**

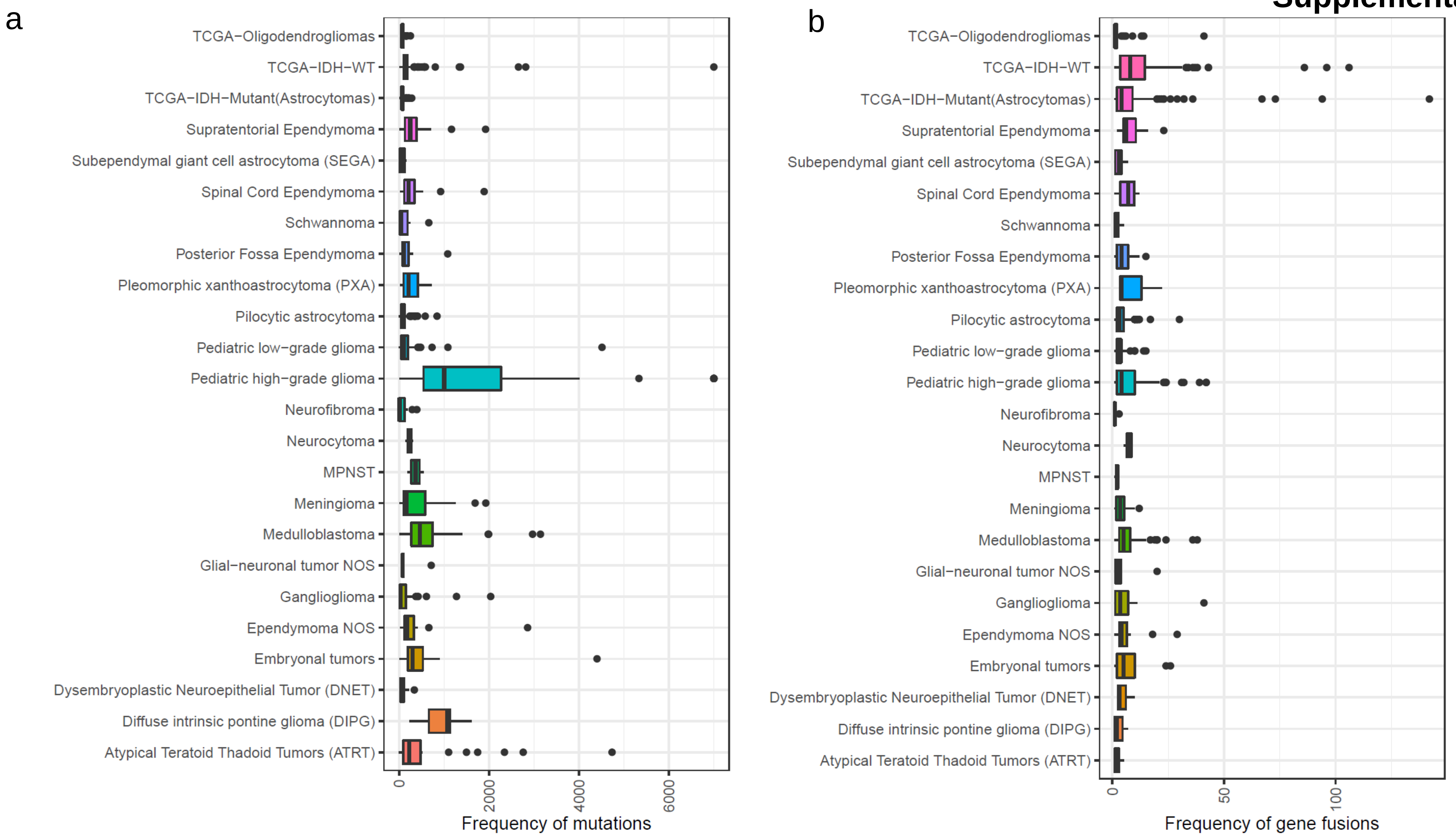


**Supplementary Figure 7.** Boxplots for adult glioma subtypes and pediatric tumor types showing frequency of (a) mutations and (b) gene fusions.

c

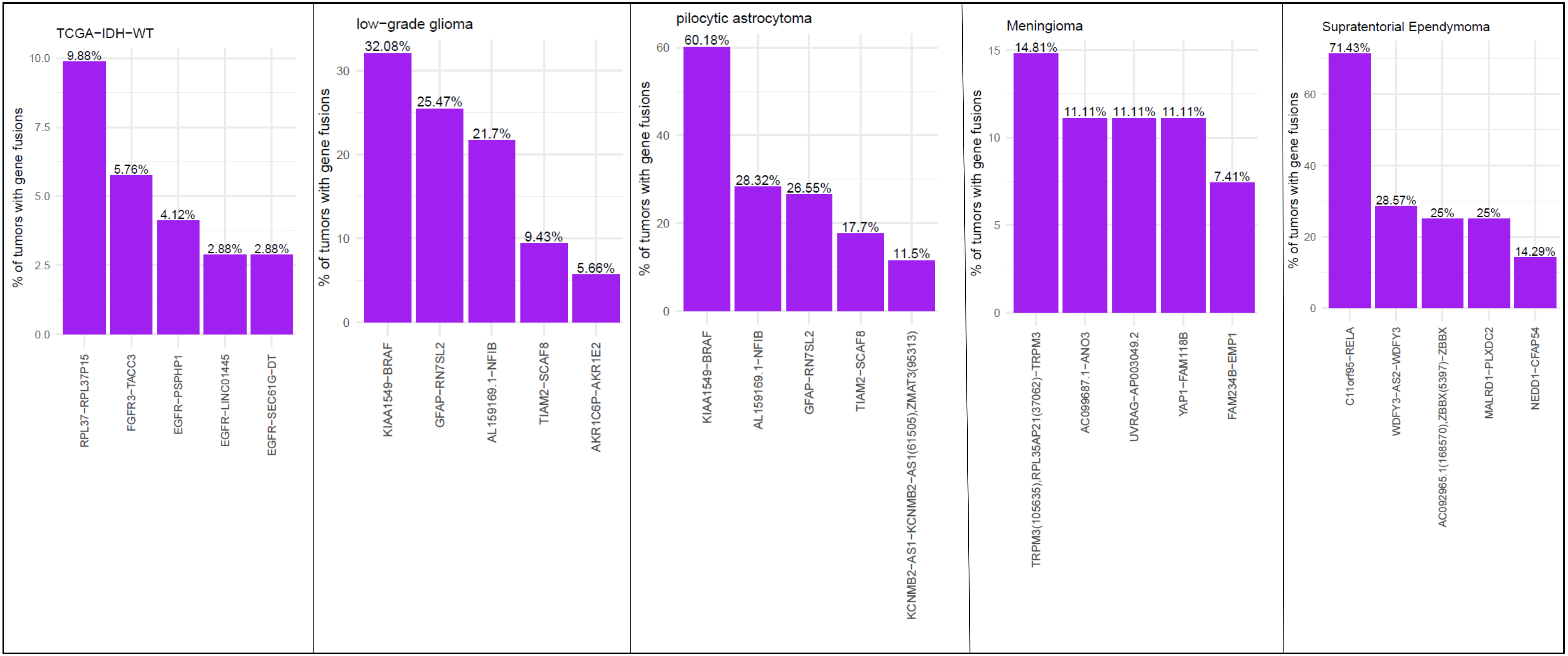


**Supplementary Figure 7.** (c) Barplots showing percentage of tumors containing top 5 gene fusions in TCGA IDH-wild type , Pediatric low-grade gliomas, pilocytic astrocytoma , meningioma and supratentorial ependymoma.

**Supplementary Fig 8a**

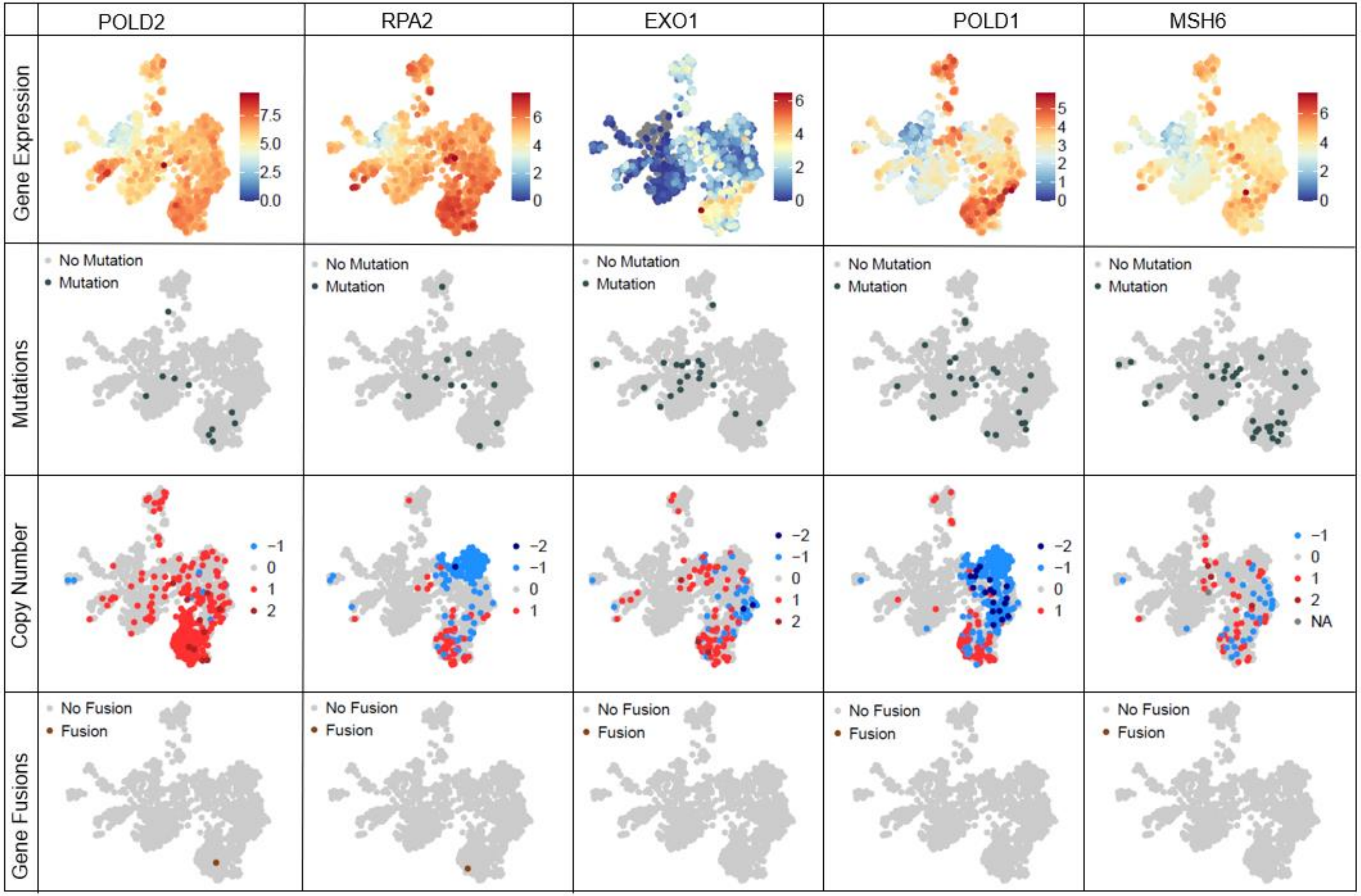


**Supplementary Figure 8 (a-c).** UMAP projection for RNASeq data from only CBTN and TCGA datasets colored in for genes from the REACTOME MISMATCH REPAIR pathway.

**Supplementary Fig 8b**

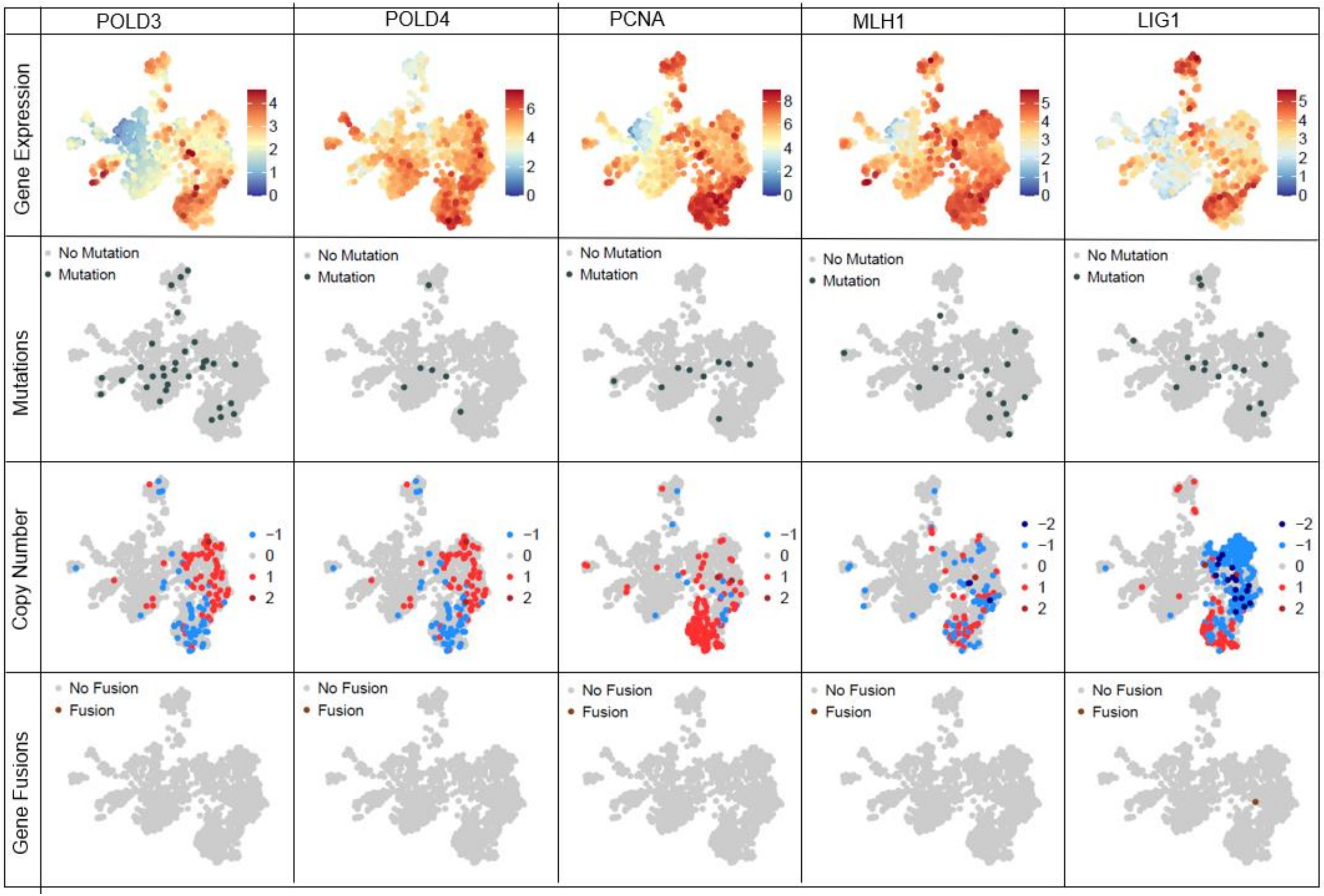


**Supplementary Figure 8 (a-c).** UMAP projection for RNASeq data from only CBTN and TCGA datasets colored in for genes from the REACTOME MISMATCH REPAIR pathway.

**Supplementary Fig 8c**

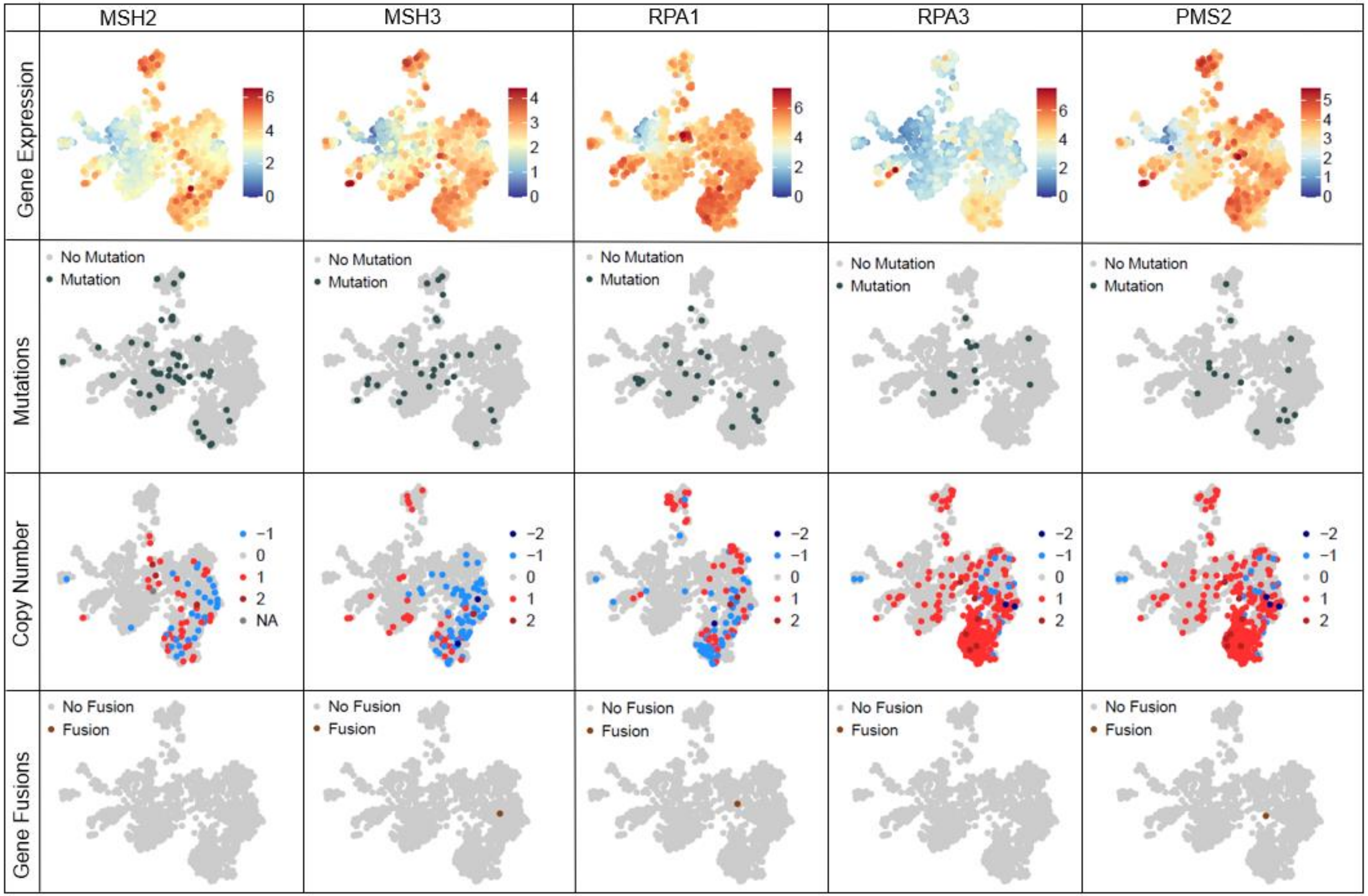


**Supplementary Figure 8 (a-c).** UMAP projection for RNASeq data from only CBTN and TCGA datasets colored in for genes from the REACTOME MISMATCH REPAIR pathway.
